# Hemogenic endothelium of the vitelline and umbilical arteries is the major contributor to mouse fetal lympho-myelopoiesis

**DOI:** 10.1101/2024.07.11.603050

**Authors:** Cristiana Barone, Giulia Quattrini, Roberto Orsenigo, Filipa Timóteo-Ferreira, Alessandro Muratore, Anna Cazzola, Arianna Patelli, Francisca Soares-da-Silva, Matthew Nicholls, Mario Mauri, Silvia Bombelli, Sofia De Marco, Deborah D’Aliberti, Silvia Spinelli, Veronica Bonalume, Alison Domingues, Gianluca Sala, Arianna Colonna, Elisabetta D’Errico, Cristina D’Orlando, Cristina Bianchi, Roberto A. Perego, Raffaella Meneveri, Marella F.T.R. De Bruijn, Ana Cumano, Alessandro Fantin, Silvia Brunelli, Rocco Piazza, Emanuele Azzoni

## Abstract

Embryonic hematopoiesis consists of distinct waves originating in rapid succession from different anatomical locations. Hematopoietic progenitors appearing earlier than definitive hematopoietic stem cells (HSCs) play key roles in fetal and postnatal life. However, their precise origin, identity and the extent of their contribution need further clarification. To this aim, we took advantage of a genetic fate-mapping strategy in mice that allows labeling and tracking of distinct subsets of hemogenic endothelium (HE). Time-course labeling of hematopoietic progenitors emerging from HE between E8.5 and E9.5, before intra-embryonic definitive HSC generation, revealed a major fetal lympho-myeloid contribution which declined in the adult. Lineage tracing coupled with whole-mount imaging and single-cell RNA sequencing located its source within hematopoietic clusters of vitelline and umbilical arteries. Functional assays confirmed the transient nature of these progenitors. We therefore unveiled a hitherto unidentified early wave of fetal-restricted hematopoietic stem/progenitor cells poised for differentiation that provide a major contribution to pre-natal hematopoiesis.

## Introduction

The vertebrate embryonic hematopoietic system is generated through consecutive and asynchronous waves of blood progenitors characterized by increasing lineage potential ^1, 2^. Despite the high degree of conservation of layered hematopoiesis ^3^, its comprehensive analysis in model organisms has historically been complicated by the fact that waves have multiple sources and overlap in time and space. Moreover, many surface markers are shared between embryonic blood progenitor cells, and their heterogeneity has only recently started to be dissected thanks to single cell methodologies ^4^.

Adult repopulating Hematopoietic Stem Cells (HSCs) are firstly and autonomously generated in the aorta-gonad-mesonephros (AGM) region starting from E10.5 in the mouse ^5, 6^. HSCs originate from a specialized population of endothelial cells termed hemogenic endothelium (HE) in the major embryonic arteries ^7-11^ and mature through a hierarchy of pro- and pre-HSC intermediates ^12, 13^. Starting from E12.0, HSCs colonize the fetal liver (FL) where they are thought to expand in numbers ^14^, and toward the end of gestation relocate to the bone marrow (BM) where they will reside throughout the length of adult life.

Prior to HSC generation and following the early onset of unipotent cells characteristic of primitive hematopoiesis, several waves of oligopotent progenitors begin emerging at E8.25 ^2, 15, 16^, initially from HE in the yolk sac (YS) ^17^. Among HSC-independent progenitors, Erythro-Myeloid Progenitors (EMPs) generate tissue resident macrophages persisting until adult life ^18-20^. EMPs were also reported to significantly contribute to fetal erythropoiesis ^21^ and fetal innate lymphoid cells ^22^. Despite the potential of EMPs to generate multiple myeloid lineages ^23^, their contribution to fetal and postnatal myelopoiesis is still unclear. Immune-restricted Lympho-Myeloid Progenitors (LMPs) emerge in the YS slightly later than EMPs and were shown to take part in fetal lympho-myelopoiesis, even though for a limited time window and contributing less than 20% of myeloid cells at E14.5 ^24^. As for fetal lymphoid cells, despite conclusive evidence that YS-derived HSC-independent B and T cell progenitors exist and persist to adulthood ^25-28^, some controversy still remains regarding the identity of the first progenitors responsible for the colonization of the fetal thymus, and in particular whether they originate from HSCs or not ^29, 30^.

HSC-independent progenitors are necessary and sufficient to sustain fetal life until the end of gestation ^31^. Indeed, recent work showed that HSCs exert a limited contribution to embryonic and fetal hematopoiesis ^32, 33^. Lineage tracing revealed the existence of embryonic multipotent progenitors (eMPPs), appearing concomitantly to definitive HSCs, that significantly contribute to fetal and adult multi-lineage hematopoiesis ^33, 34^. Although the origin of eMPPs appears to be HSC-independent ^33, 35, 36^, it is not clear when and where they emerge during development. The existence of fetal HSCs with characteristics distinct from adult HSCs has also been suggested ^37, 38^, however, as for eMPPs, a prospective identification of these progenitors, which would allow localization in their niche of emergence, is currently not possible. Moreover, the true extent of their contribution to fetal and adult hematopoiesis needs further clarification.

To genetically label and trace discrete subsets of intra- and extra-embryonic HE, we took advantage of well-established conditional fate-mapping strategies in mouse. We found that a wave of fetal-restricted hematopoietic stem/progenitor cells (HSPCs) emerges from HE between E8.5 and E9.5, before the onset of adult-type HSCs, and acts as the main driver of fetal lympho-myelopoiesis. Through a combination of whole-mount imaging, single cell transcriptomics and functional assays, we pinpointed the initial emergence of HSPCs belonging to this wave to pre-HSCs localized within hematopoietic clusters of the vitelline and umbilical arteries (VU). Moreover, we show that fetal-restricted HSPCs are not endowed with long-term multilineage engraftment potential, but instead represent a subset of hematopoietic progenitors poised for differentiation. We hypothesize that eMPPs and fetal HSCs are generated within this hematopoietic wave.

## Results

### HE lineage tracing identifies a population of fetal-restricted hematopoietic progenitors

All hematopoietic cells other than primitive erythrocytes derive from *Cdh5*+ HE ^1^. To differentially label embryonic and fetal hematopoietic waves and systematically trace their contribution during fetal and adult hematopoiesis, we employed a well validated pulse-chase approach driven by tamoxifen-inducible *Cdh5-CreER^T2^* ^39^, in combination with suitable reporter lines (*R26^tdTomato^*, *R26^zsGreen^*or *R26^EYFP^*) (**Figure S1A**). An advantage of this strategy is to use just one Cre mouse line, thus avoiding bias from cell type-specific promoters ^40^. As Tamoxifen metabolization into its active metabolite 4-hydroxytamoxifen (4-OHT) takes 6 to 12 hours *in vivo* in mice and is still detected in the serum at 48 hours post-administration ^41^, we directly used (Z)-4-OHT, which has a *in vivo* half-life of <3 hours, peaks in the serum almost immediately after administration and is largely cleared to undetectable levels within a 12-hour window as recently shown by mass spectrometry ^42^.

We first evaluated the labeling of YS EMPs (Ter119-Kit+ CD41^low^ CD16/32+)^23^. In *Cdh5-CreER^T2^::R26^zsGreen^* embryos, E9.5 and E10.5 YS EMPs were found labeled with high efficiency with 4-OHT activation at both E7.5 and E8.5 (**Figure S1B,C**). EMP labeling was also confirmed by whole-mount imaging of E9.5 YS (**Figure S1D**), which identified no significant differences in the number of labeled Kit+ clusters when traced at the two activation time points (**Figure S1E**). Consistent with this, brain microglia, which originates from YS EMPs ^43^, was found highly labeled with both 4-OHT at E7.5 and E8.5, in the E16.5 fetus and in the adult (**Figure S1F,G**). In contrast, as expected, 4-OHT at E10.5 did not label fetal or adult microglia (**Figure S1G**).

We analyzed the labeling of LMPs (Lin-Kit+ CD45+ Flt3+ IL7Rα+) in *Cdh5-CreER^T2^::R26^tdTomato^*E11.5 FL. With 4-OHT at 7.5, the observed recombination frequency was low in LMPs and only became partial (30%) with activation at E8.5 (**Figure S1H,I**), consistent with LMP emergence at/around E9.5 in the YS ^24^. Interestingly, the percentage of traced CD45+ Kit+ hematopoietic progenitors in the E11.5 FL doubled when 4-OHT was delivered at E8.5 compared to E7.5 (**Figure S1J**), suggesting that, at least in part, they may originate independently from EMPs.

Next, we investigated the extent of labeling within phenotypic pre-HSCs type I (CD41^low^ CD45-CD43+) and II (CD41^low^ CD45+ CD43+) in both AGM and YS (including VU) of *Cdh5-CreER^T2^::R26^tdTomato/zsGreen^*mice at E11.5 (**Figure 1A**). Strikingly, the significant majority of phenotypic pre-HSCs at this stage was labeled with 4-OHT at E8.5, whereas only a minor fraction traced with 4-OHT at E10.5 (**Figure 1B-C**).

**Figure 1.**
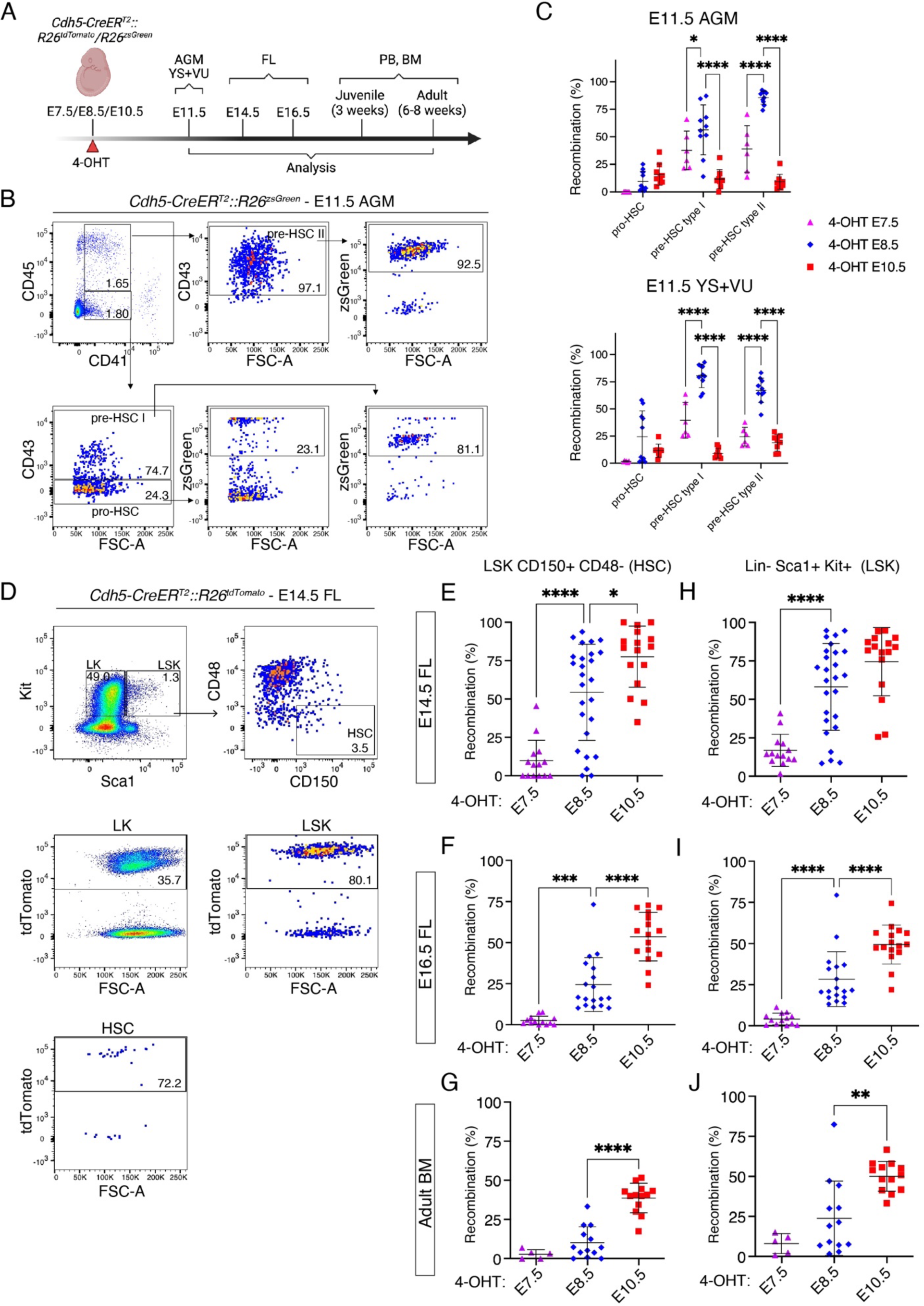
Lineage tracing of HE identifies fetal-restricted HSPCs. **(A)** Visual schematic of lineage tracing experiments in *Cdh5-CreER^T2^::R26^tdTomato^/ R26^zsGreen^* E11.5-E14.5-E16.5 embryos and postnatal mice. **(B)** Representative flow cytometric analysis of aorta-gonad-mesonephros (AGM) region pro-HSCs (CD45-CD41+ CD43-), pre-HSCs type I (CD45-CD41+ CD43+) and pre-HSCs type II (CD45+ CD41+ CD43+) in E11.5 *Cdh5-CreER^T2^::R26^zsGreen^* embryos. 4-OHT at E7.5 (n = 6), 4-OHT at E8.5 (n = 10), 4-OHT at E10.5 (n = 8) Yolk Sac + Vitelline and Umbilical artery (YS+VU) and 4-OHT at E7.5 (n = 6), 4-OHT at E8.5 (n = 10), shown here, 4-OHT at E10.5 (n = 9) AGM were analyzed individually in 3 independent experiments. **(C)** Quantification of flow cytometric analysis displayed in **(B)**. The percentage (%) of recombination is represented as the % of zsGreen+ or tdTomato+ cells within pro/pre-HSCs. Error bars represent mean ± standard deviation (SD). *p<0.05, ****p<0.0001 (two-way ANOVA followed by Tukey’s multiple comparisons test). **(D)** Representative flow cytometric analysis of fetal liver (FL) HSCs (Lineage-Kit+ Sca1+ CD48-CD150+), LK (Lineage-Kit+ Sca1-) and LSK (Lineage-Kit+ Sca1+) and the relative percentage of labeling in E14.5 *Cdh5-CreER^T2^::R26^tdTomato^*embryos. Lineage cocktail: B220, CD19, CD3e, F4/80, Gr1, Nk1.1, Ter119, 7-AAD. 4-OHT at E7.5 (n = 14), 4-OHT at E8.5 (n = 26), 4-OHT at E10.5 (n = 16), shown here, FL were analyzed individually in 6 independent experiments. (**E-G**) Quantification of flow cytometric analysis displayed in **(D)**. The % of recombination is represented as the % of tdTomato+ or zsGreen+ cells within HSCs in E14.5 FL (**E**), E16.5 FL (**F**) or adult bone marrow (BM) (**G**). 4-OHT at E7.5 (n = 13), 4-OHT at E8.5 (n = 18) and 4-OHT at E10.5 (n = 17) E16.5 FL were analyzed individually in 4 independent experiments. 4-OHT at E7.5 (n = 5), 4-OHT at E8.5 (n = 13), 4-OHT at E10.5 (n = 13) BM were analyzed individually in 7 independent experiments. Error bars represent mean ± standard deviation (SD). *:p<0.05, ***:p<0.001, ****:p<0.0001 (one-way ANOVA followed by Tukey’s multiple comparisons test). (**H-J**) Quantification of flow cytometric analysis displayed in **(D)**. The % of recombination is represented as the % of tdTomato+ or zsGreen+ cells within LSK in E14.5 FL (**H**), E16.5 FL (**I**) or adult bone marrow (**J**) (BM). Replicates as indicated in D-G. Error bars represent mean ± standard deviation (SD). **:p<0.01, ****:p<0.0001 (one-way ANOVA followed by Tukey’s multiple comparisons test).

We then tested the labeling of fetal and adult phenotypic HSCs (Lin-Kit+ Sca1+ CD150+ CD48-) (**Figure 1D**). E14.5, E16.5 FL and adult bone marrow (BM) HSCs were extensively labeled with 4-OHT at E10.5, but not at E7.5 (**Figure 1E-G**), confirming a previous report ^44^. Interestingly, 4-OHT at E8.5 labeled E14.5 FL HSCs with variable efficiency (58% on average; **Figure 1E**) not dependent on the specific reporter line (**Figure S1K**), but decreasing to an average of 25% at E16.5 (**Figure 1F**) and 10% in the adult BM (**Figure 1G**), raising the possibility of a wave of fetal-restricted HSCs as previously suggested ^37^. A similar labeling pattern, however, was also observed in Lin-Kit+ Sca1+ (LSK) progenitors (**Figure 1H-J**). Non-HSC hematopoietic progenitors (LK, Lin-Kit+ Sca1-), comprising granulocyte-macrophage progenitors (GMP), common myeloid progenitors (CMP) and megakaryocyte-erythroid progenitors (MEP) labeling was at the highest with 4-OHT at E8.5 in E14.5 FL (**Figure S1L**), whereas at E16.5 and in adult BM they were mostly labeled with 4-OHT at E10.5 (**Figure S1M-N**).

Taken together, these data show that HE lineage tracing with 4-OHT activation at E8.5 in the *Cdh5-CreER^T2^* model phenotypically identifies a putative population of fetal-restricted HSPCs. In contrast, 4-OHT pulses at E7.5 or E10.5 respectively label either EMPs or phenotypic adult-type HSCs. Therefore, the same genetic model can be used to study the relative fetal and adult contributions of three sequential waves of HE in an unbiased way. Importantly, differential labeling of immunophenotypic pre-HSCs at E11.5 suggests that, at least until this stage, their majority represent precursors of fetal-restricted HSPCs, whereas the pre-HSCs that give rise to adult-type HSCs likely have yet to emerge or mature.

### Fetal lympho-myelopoiesis is largely derived from hematopoietic progenitors emerging from HE between E8.5 and E9.5

To examine the fetal lympho-myeloid contribution of the three waves of HE-derived hematopoietic precursors at E7.5, E8.5 and E10.5, we analyzed those lineages in *Cdh5-CreER^T2^::R26^tdTomato^* E16.5 FL and thymus. 4-OHT activation at E7.5 yielded an average labeling of only 35% of F4/80+ CD11b^low^ macrophages, less than 10% B cells, 10-15% T cells and 5% F4/80^low/-^ CD11b+ myeloid cells (**Figure 2A,B; Figure S2A,B**) in the E16.5 FL and thymus, suggesting that EMPs do not significantly contribute to fetal lympho-myelopoiesis. E10.5 activation resulted in labeling of 30% of B and myeloid cells (**Figure 2A; Figure S2A**), a similar contribution to T cells, with higher labeling (30-40%) in less differentiated thymocytes (DN1 and DN2) (**Figure 2B; Figure S2B**) and negligible labeling in macrophages. Strikingly, the highest labeling frequencies in all lineages were observed with 4-OHT activation at E8.5, yielding on average 70% of labeled B and myeloid cells in the E16.5 FL (**Figure 2A; Figure S2A**) and 50-60% of labeled E16.5 fetal thymocytes (**Figure 2B; Figure S2B**). In contrast to E10.5 activation, double-positive (CD4+ CD8+ DP) T cells showed the highest labeling frequency, and embryonic ψο T cells were almost exclusively labeled by 4-OHT at E8.5 (**Figure 2B**), suggesting that our tracing system can differentially label the two separate waves of thymic-settling progenitors (TSP) ^45^.

**Figure 2.**
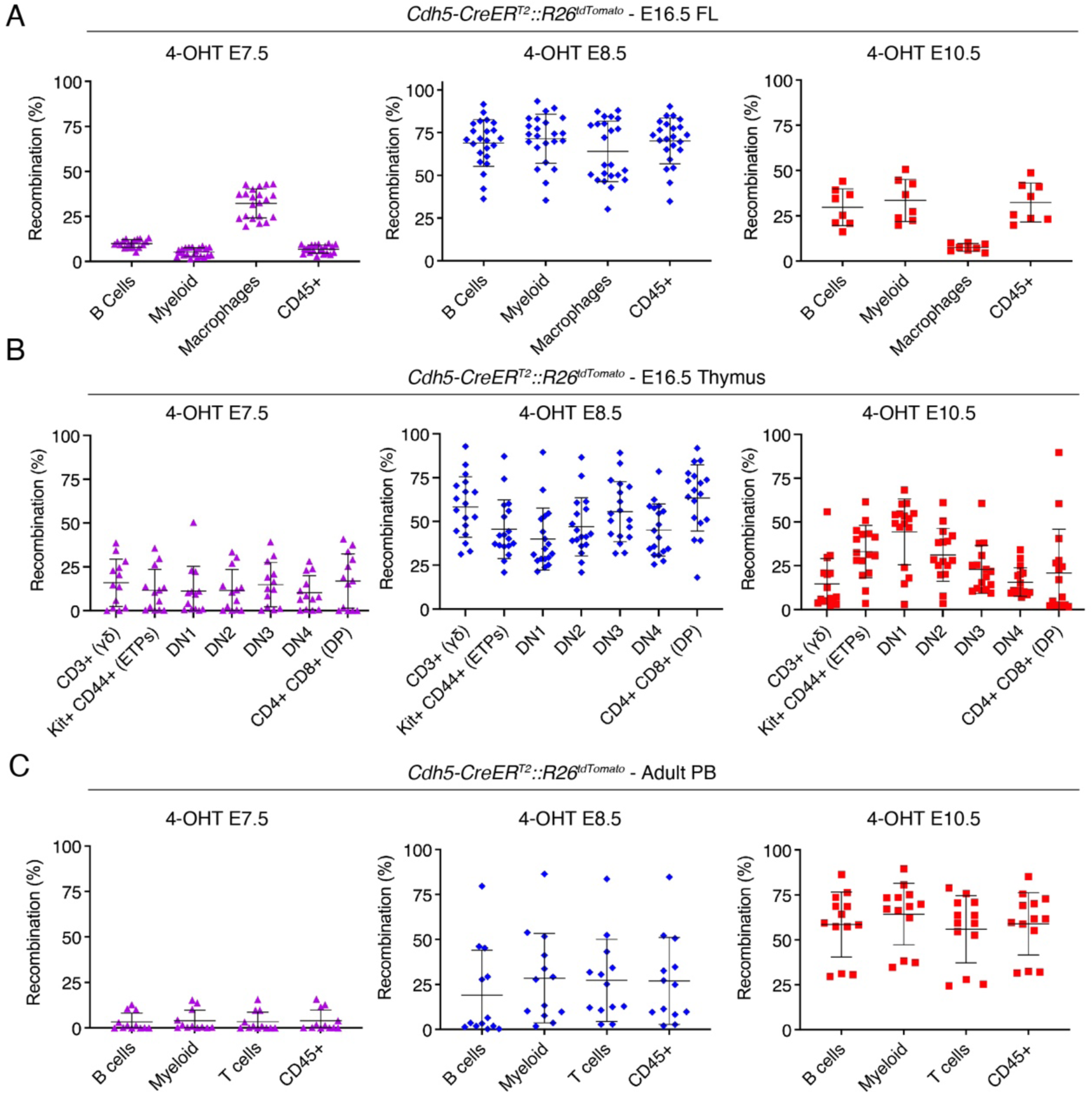
Extensive lympho-myeloid contribution of fetal-restricted HSPCs at the end of gestation. **(A)** Quantification of flow cytometric analysis of labeled B cells (CD45+ B220+), myeloid cells (CD45+ CD11b+), macrophages (CD45+ F4/80^hi^) and CD45+ cells (leukocytes) in *Cdh5-CreER^T2^::R26^tdTomato^* E16.5 FL activated with 4-OHT at E7.5 (left), E8.5 (middle) or E10.5 (right). 4-OHT E7.5 (n = 21), 4-OHT E8.5 (n = 23), N = 8 (4-OHT E10.5) FL were analyzed individually in 5 independent experiments (gating strategy in S2A). Error bars represent mean ± SD. **(B)** Quantification of flow cytometric analysis of labeled thymocytes in *Cdh5-CreER^T2^::R26^tdTomato^* E16.5 fetal thymus (gating strategy in S2B), activated with 4-OHT at E7.5 (left), E8.5 (middle) or E10.5 (right). 4-OHT E7.5 (n = 13), 4-OHT E8.5 (n = 18), or 4-OHT E10.5 (n = 16) thymuses were analyzed individually in 4 independent experiments. Error bars represent mean ± SD. **(C)** Quantification of flow cytometric analysis of labeled B cells (CD45+ B220+), myeloid (CD45+ CD11b+), T cells (CD45+ CD3e+) and CD45+ cells (leukocytes) in adult (2 months old) PB from *Cdh5-CreER^T2^::R26^tdTomato^* mice (gating strategy in S2D), activated with 4-OHT at E7.5 (left), E8.5 (middle) or E10.5 (right). 4-OHT E7.5 (n = 12), 4-OHT E8.5 (n = 13), 4-OHT E10.5 (n = 13) mice were analyzed individually in 7 independent experiments. Error bars represent mean ± SD.

Post-natal analysis confirmed absence of peripheral blood (PB) labeling (<5%) in all lineages with activation at E7.5, both at 21 days (**Figure S2C**) and 2 months (**Figure 2C; Figure S2D**). 4-OHT at E10.5 resulted in high levels of recombination in all lineages gradually increasing between 21 days and 2 months (**Figure 2C; Figure S2C**), reflecting the transition to adult-type definitive HSC-derived hematopoiesis ^46, 47^. Conversely, activation at E8.5 showed highly variable labeling at 21 days (average 30-35%) (**Figure S2C**), which dropped to 20-25% at 2 months (**Figure 2C**).

These data show that the hematopoietic wave that emerges from HE between E8.5 and E9.5 is the main contributor of fetal lympho-myelopoiesis. As similar labeling dynamics were seen with the same activation mode for LK, LSK and phenotypic HSC (**Figure 1, Figure S1**), this wave of HE likely contains the precursors that generate a pre-constituted hierarchy of fetal-restricted progenitors, including fetal HSCs, HSC-independent eMPPs and other progenitors ^33, 34, 48, 49^. We will collectively refer to these as “fetal-restricted HSPCs”.

### Fetal-restricted HSPCs first emerge from HE of the vitelline and umbilical arteries

To get insight into the dynamics of fetal-restricted HSPC generation, we investigated sites of hematopoietic emergence using whole-mount confocal imaging. As mentioned, Kit+ CD31+ hematopoietic clusters in the E9.5 YS, corresponding to EMPs, were equally labeled by 4-OHT activation at E7.5 or E8.5 (**Figure S1D,E**). CD31+ Kit+ hematopoietic clusters in the dorsal aorta (DA), thought to contain pre-HSCs ^50^, peak at E10.5 ^51^. However, the first intra-embryonic Kit+ hematopoietic clusters do not appear within the DA, but in the vitelline artery (VA) at E9.5 ^12, 52, 53^. Although the majority of these clusters are thought to contain progenitor cells other than pro-HSC ^12^, major extra-embryonic arteries are known to represent sites of pre-HSC emergence ^10, 54^. Importantly, because of the lack of specific ways to trace these early progenitors, their contribution has never been analyzed in an unperturbed system. Remarkably, few Kit+ hematopoietic clusters in the *Cdh5-CreER^T2^::R26^EYFP^*E9.5 VA were labeled by 4-OHT at E7.5, but, in contrast, their majority was labeled by the E8.5 activation (**Figure 3A,B**), correlating with the emergence of fetal-restricted HSPCs. These observations indicate that such progenitors first emerge within the VA. Flow cytometry analysis of E9.5 hematopoietic progenitors other than EMP (non-EMPs, Kit+ CD41^low^ CD16/32-) showed absence of differential labeling in the YS, but significantly higher labeling within the caudal part of the embryo (CP) when traced at E8.5 (**Figure S3A**), consistent with imaging data and an identity of VA clusters independent from EMPs. Next, we evaluated labeling of hematopoietic clusters in the VA and umbilical artery (UA) of E10.5 embryos and compared it to the DA. 4-OHT activation at E7.5 yielded low labeling frequency of both VU and aortic clusters (**Figure 3C,D**). Conversely, 4-OHT at E8.5 resulted in high levels of recombination in VU clusters, but significantly lower in DA clusters (**Figure 3C,D**).

**Figure 3.**
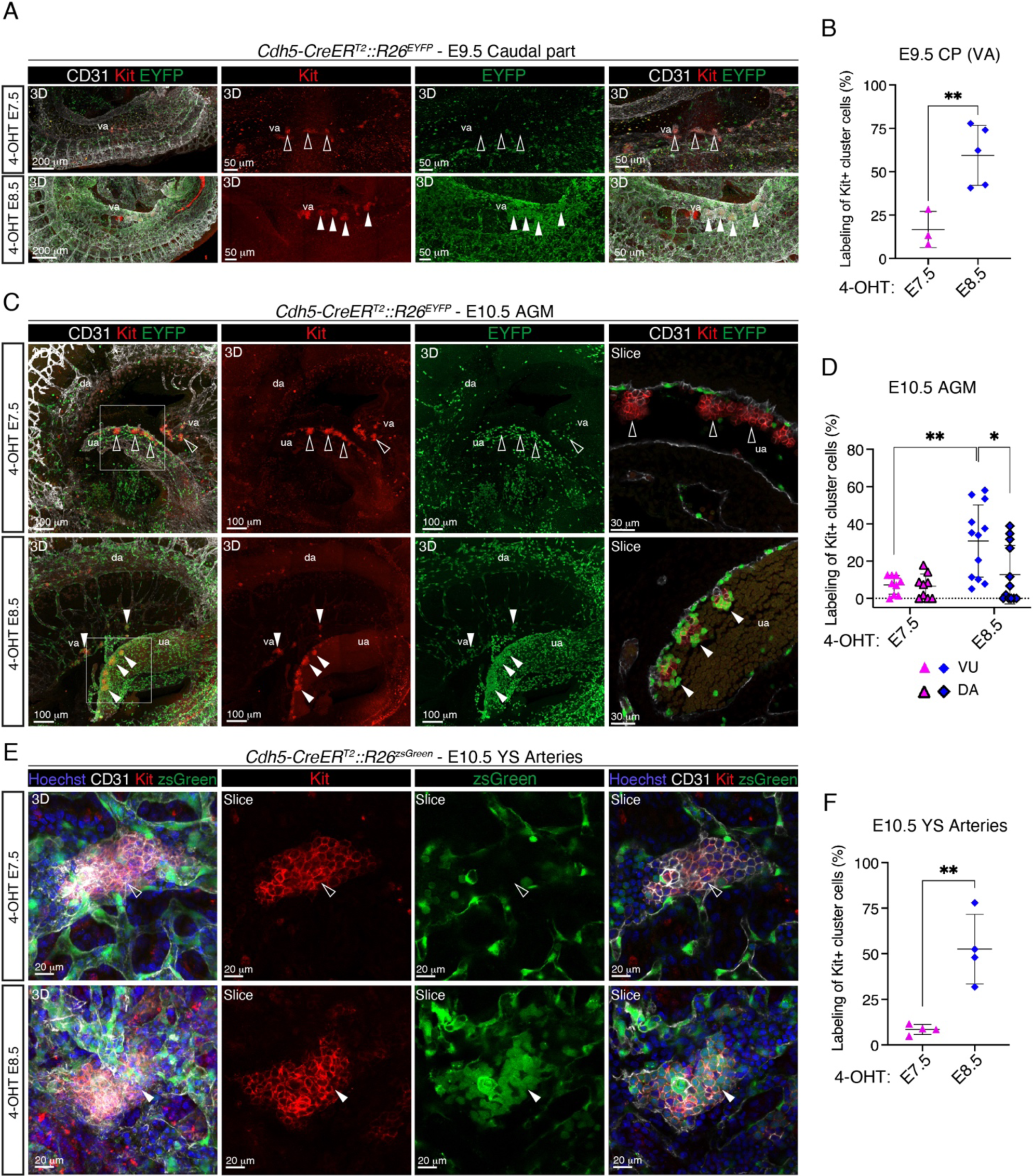
4-OHT activation at E8.5 in *Cdh5-CreER^T2^* embryos preferentially labels hematopoietic clusters in vitelline and umbilical arteries. **(A)** Confocal whole mount immunofluorescence (WM-IF) analysis of E9.5 (21-27sp) *Cdh5-CreER^T2^::R26^EYFP^*embryos within the caudal part (CP). Images show maximum intensity 3D projections. Middle and right panels are magnified images. Arrowheads indicate Kit+ hematopoietic clusters in the vitelline artery (va), labeled with 4-OHT at E8.5 (white), but not 4-OHT at E7.5 (empty). Scale bars: 200 μm (left), 50 μm (magnifications). **(B)** Labeling quantification of Kit+ cluster cells in the VA of *Cdh5-CreER^T2^::R26^EYFP^* E9.5 embryos as shown in **(A)**. Measurements were performed on images from 4-OHT E7.5 (n=3), and E8.5 (n=5) different embryos (1-4 images / embryo); 11 (4-OHT E7.5), 6 (4-OHT E8.5) different images used. Error bars represent mean ± SD. **:p<0.01 (two-tailed unpaired Student’s *t*-test). **(C)** Confocal WM-IF analysis of AGM region of E10.5 (32-36sp) *Cdh5-CreER^T2^::R26^EYFP^* embryos. Left and middle panels show maximum intensity 3D projections. Boxed area in the merged image is magnified in the right panel and shows a single 2.5 μm-thick optical slice. Arrowheads indicate Kit+ hematopoietic clusters in the umbilical artery (ua), unlabeled with 4-OHT at E7.5 (empty arrowheads; top), or labeled at E8.5 (white arrowheads; bottom). 4-OHT E7.5 (n = 9), and 4-OHT E8.5 (n = 12) different embryos were analyzed in 7 independent experiments. Scale bars: 100 μm (3D), 30 μm (slice). va: vitelline artery; da: dorsal aorta. **(D)** Labeling quantification of Kit+ hematopoietic cluster cells in the AGM of E10.5 *Cdh5-CreER^T2^::R26^EYFP^*embryos as displayed in **(C)**. Dorsal aorta (DA) clusters were quantified separately from clusters in vitelline and umbilical arteries (VU). Measurements were performed on images from 4-OHT E7.5 (n = 9), and 4-OHT E8.5 (n = 12) different embryos (3-6 images / embryo); 39 (4-OHT E7.5), 56 (4-OHT E8.5) different images used. Error bars represent mean ± SD. *:p<0.05, **:p<0.01 (two-way ANOVA followed by Tukey’s multiple comparisons test). **(E)** Confocal WM-IF analysis of E10.5 *Cdh5-CreER^T2^::R26^zsGreen^*YS large arteries. Left panels show 3D maximum intensity projections. Middle and right panels show single 2.5 μm-thick optical slices. Arrowheads indicate large Kit+ hematopoietic clusters, unlabeled with 4-OHT at E7.5 (empty arrowheads; top), or labeled at E8.5 (white arrowheads; bottom). 4-OHT E7.5 (n=4), and 4-OHT E8.5 (n=4) different YS were analyzed in 2 independent experiments. Scale bars: 20 μm. **(F)** Labeling quantification of Kit+ cluster cells in the YS large arteries as displayed in **(E)**. Measurements were performed on images from 4-OHT E7.5 (n=4), and 4-OHT E8.5 (n=4) different YSs (2-4 images / YS); 12 (4-OHT E7.5), 12 (4-OHT E8.5) different images used. Error bars represent mean ± SD. **:p<0.01 (two-tailed unpaired Student’s *t*-test).

Previous *ex vivo* culture assays revealed the presence of pre-HSCs in the YS at E10.5, but not at E9.5 ^12, 52, 55^; from E11.5, YS pre-HSCs seem to decline ^13, 14, 55^. Whole-mount analysis of E10.5 YS identified the presence of large Kit+ hematopoietic clusters, which were absent at E9.5. These clusters exclusively localized in the VA and its ramifications, and were previously documented to express Ly6a ^53^ and Hlf ^56^, markers associated with HSC activity. The majority of cells in these clusters in *Cdh5-CreER^T2^::R26^zsGreen^* YS were not labeled with activation at E7.5 but, instead, were consistently labeled with 4-OHT at E8.5 (**Figure 3E,F**). In line with these results, differential labeling of non-EMPs was observed in both YS and AGM at E10.5 (**Figure S3B**).

Overall, these results suggest that fetal-restricted HSPCs first emerge from the HE of the VA and the UA. At E9.5, the location of the hematopoietic clusters likely to contain these progenitors is solely intra-embryonic, whereas from E10.5 is both intra- and extra-embryonic, still maintaining a close association with the VU.

### EMPs do not significantly contribute to hematopoietic clusters in intra- and extra-embryonic arteries

In order to gain insight on the origin of hematopoietic clusters in different sites of emergence, we performed lineage tracing using the *Csf1r-iCre* transgenic mouse line, which targets EMPs and their progeny ^18, 19, 40, 57^.

Kit+ clusters in the E9.5 VA of *Csf1r-iCre::R26^tdTomato^*embryos showed near complete absence of labeling (**Figure 4A**), despite the expected highly efficient labeling of E9.5 EMPs (**Figure 4B**). Non-EMPs (Kit+ CD41^low^ CD16/32-) were not labeled in the E9.5 CP, whereas ∼50% recombined in the YS, possibly representing immature EMPs (**Figure 4B**). Hematopoietic clusters in the E10.5 DA, VA, UA and large arterial clusters in the E10.5 YS showed very low labeling frequencies, confirming that they do not originate from EMPs (**Figure 4C,D**). In contrast, extensive labeling was detected in the E10.5 FL, in agreement with its early colonization by EMPs (**Figure 4C,D**). Accordingly, at E10.5 EMP labeling remained high (>90%), whereas less than 25% non-EMPs were labeled in both YS+VU and embryo proper (**Figure 4E**). E9.5-E10.5 erythroid cells were not labeled, and megakaryocytes (Mk) showed low recombination rates at E9.5, increasing to ∼50% at E10.5 (**Figure 4B,E**). These observations are consistent with a previous report that employed an inducible Cre line driven by the *Csf1r* promoter ^58^, thus reinforcing the notion that these lineages initially derive from direct *in situ* differentiation of unipotent progenitors other than *Csf1r*+ EMPs. These data confirm that hematopoietic clusters emerging in the major intra- and extra-embryonic arteries, including the E10.5 YS, largely contain cells that develop independently of EMPs.

**Figure 4.**
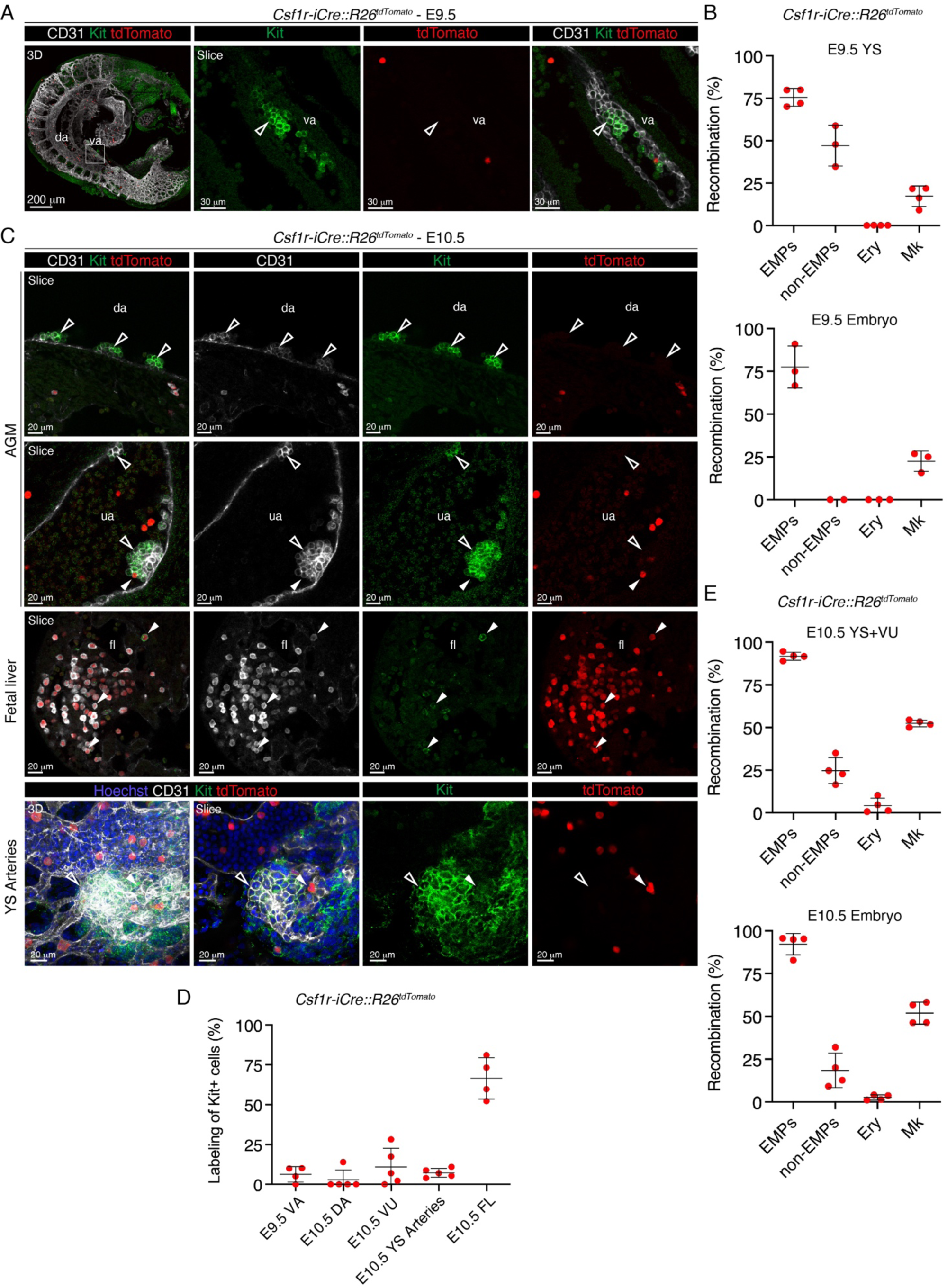
*Csf1r-iCre* lineage tracing reveals a non-EMP identity of hematopoietic clusters in intra- and extra-embryonic arteries. **(A)** Confocal WM-IF analysis of E9.5 *Csf1r-iCre::R26^tdTomato^* embryos. 3D magnification (left panel) or single 2.5 μm-thick optical slices (middle and right panels) are shown. Empty arrowheads indicate unlabeled (tdTomato-) Kit+ clusters in the va. 4 Embryos (n=4) were analyzed in 2 independent experiments. Scale bar: 200 μm (3D), 30 μm (slices). **(B)** Quantification of flow cytometric analysis of labeled (tdTomato+) EMPs (Ter119-Kit+ CD41^lo^ CD16/32+), non-EMPs (Ter119-Kit+ CD41^lo^ CD16/32-), erythrocytes (Ter119+) and megakaryocytes (Ter119-Kit-CD41^hi^) in E9.5 *Csf1r-iCre::R26^tdTomato^* YSs (top) and embryos (bottom). Gates as shown in Figure S1B. A total number of 4 different YS and 4 embryos were analyzed. Error bars represent mean ± SD. **(C)** Confocal WM-IF analysis of E10.5 *Csf1r-iCre::R26^tdTomato^* AGM region (top two rows), FL (middle) and YS large arteries (bottom). Left YS panel shows a 3D maximum intensity projection; other images are single 2.5 μm-thick optical slices. Arrowheads indicate Kit+ hematopoietic clusters in the da, ua and hematopoietic cells in the FL. Empty arrowheads indicate unlabeled (tdTomato-) cells, white arrowheads indicate labeled (tdTomato+) ones. A total number of 5 different embryos were analyzed in 3 independent experiments. **(D)** Quantification of WM-IF analysis in **(C)**. Measurements were performed on images from different E9.5 (n=4) and E10.5 embryos (n=5) (2-8 images/embryo) in 3 independent experiments; 21 (E9.5) and 40 (E10.5) different images used. Error bars represent mean ± SD. **(E)** Quantification of flow cytometric analysis of labeled (tdTomato+) EMPs (Ter119-Kit+ CD41^lo^ CD16/32+), non-EMPs (Ter119-Kit+ CD41^lo^ CD16/32-), erythrocytes (Ter119+) and megakaryocytes (Ter119-Kit-CD41^hi^) in E10.5 *Csf1r-iCre::R26^tdTomato^* YS+VU (top) and embryos (bottom). A total number of 4 different YS, and 4 embryos were analyzed. Error bars represent mean ± SD.

### B- and T-lymphoid potential appears in intra- and extra-embryonic HE between E8.5 and E9.5

To determine whether the *ex vivo* potential of fetal-restricted HSPCs mirrored their *in vivo* fate, we isolated traced and untraced cells from *Cdh5-CreER^T2^::R26^tdTomato/zsGreen^*E9.5 YS or CP and E10.5 YS including the extra-embryonic portion of the VA and UA (YS+VU) or AGM (**Figure 5A,B**). We performed colony-forming unit-culture (CFU-C) assays able to detect single or combined erythroid, myeloid and Mk potential of mature progenitors. CFU-Cs almost completely segregated with labeled cells at both activations in either YS or CP/AGM (**Figure 5C**), in agreement with the absence of differential labeling of EMPs at these two time points (**Figure S1B-E**).

**Figure 5.**
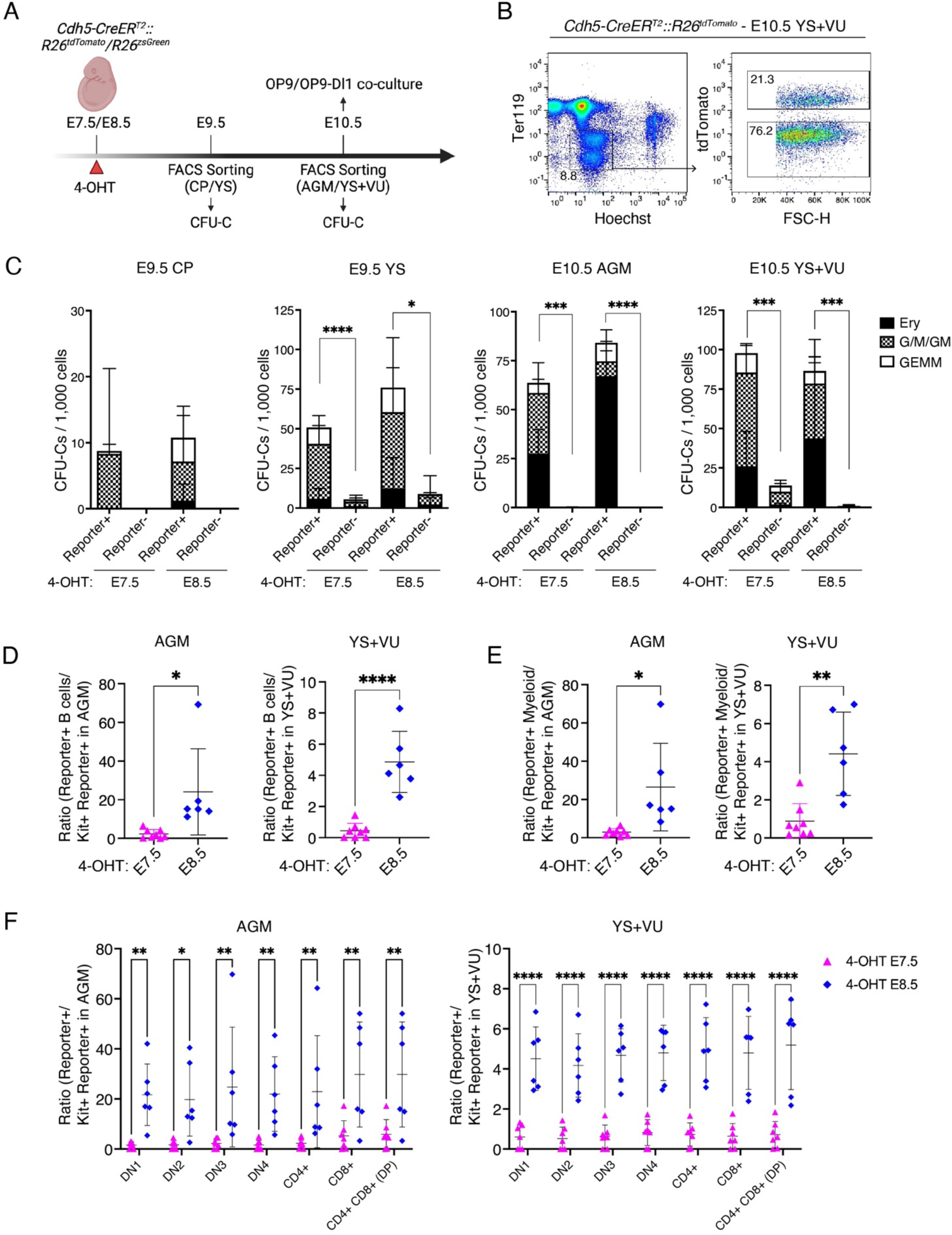
*Ex vivo* B and T lymphoid potential emerges in intra- and extra-embryonic HE at E8.5, while erythro-myeloid potential is already present at E7.5. **(A)** Visual schematic of *ex vivo* Colony-forming unit-culture (CFU-C) and OP9 co-culture experiments. **(B)** Representative flow cytometric gating strategy for the isolation of live Ter119-labeled (tdTomato+ or zsGreen+) and unlabeled (tdTomato- or zsGreen-) cells for CFU-C assays from E10.5 *Cdh5-CreER^T2^*YS+VU, activated with 4-OHT at E7.5 or E8.5 (shown here). **(C)** CFU-C numbers obtained from labeled or unlabeled live Ter119-cell fractions from *Cdh5-CreER^T2^::R26^tdTomato^*or *Cdh5-CreER^T2^::R26^zsGreen^* E9.5 Caudal part (CP) and YS (left) or E10.5 AGM and YS+VU (right), activated with 4-OHT at E7.5 or E8.5. Numbers of colonies are normalized to 1,000 cells seeded. E9.5 4-OHT E7.5 (n=12), E9.5 4-OHT E8.5 (n=6), E10.5 4-OHT E7.5 (n=4), E10.5 4-OHT E8.5 (n=4) different samples were analyzed in 6 independent experiments. Starting number of YS cells seeded: 100-4,300. Starting number of CP or AGM cells seeded: 40-150,000. GEMM: granulocyte, erythroid, monocyte/macrophage, megakaryocyte; G/M/GM: granulocyte, monocyte/macrophage; Ery: erythroid. Data presented as mean ± SD. *:p<0.05, ***:p<0.001, ****:p<0.0001, E9.5 CP 4-OHT E7.5: p=0.1204, E9.5 CP 4-OHT E8.5: p= 0.0988 (two-tailed unpaired Student’s *t*-test). **(D-E)** OP9 (B cell) co-culture assays. Values represent the frequency of labeled B (CD45+ AA4.1+ B220+ CD19+) **(D)** and myeloid (CD45+ AA4.1-CD11b+) **(E)** cells at day (d)10 of co-culture, normalized to the percentage of labeled Kit+ cells at d0 of co-culture, from E10.5 AGM and YS+VU activated with 4-OHT at E7.5 or E8.5. Representative flow cytometric analysis of OP9 co-cultures is shown in Figure S3D. 4-OHT E7.5 B cells (n = 8), 4-OHT E7.5 myeloid cells (n = 8), 4-OHT E8.5 B cells (n = 6), and 4-OHT E8.5 myeloid cells (n = 6 different samples were analyzed in 4 independent experiments. Error bars represent mean ± SD. *:p<0.05, ***:p<0.001, ****:p<0.000 (two-tailed unpaired Student’s *t*-test). **(F)** OP9-Dl1 (T cell) co-culture assays. Values represent the frequency of labeled T cells at day (d)10 of co-culture normalized on the percentage of labeled Kit+ cells at d0 of co-culture, from E10.5 AGM and YS+VU activated with 4-OHT at E7.5 or E8.5. Representative flow cytometric analysis of OP9-Dl1 co-cultures is shown in Figure S3E. 4-OHT E7.5 (n = 7) and 4-OHT E8.5 (n = 6) different samples were analyzed in 4 independent experiments. Error bars represent mean ± SD. *:p<0.05, **:p<0.01, ****:p<0.0001 (two-way ANOVA followed by Tukey’s multiple comparisons test).

We next tested B- and T-lymphoid potential of HE using OP9 and OP9-DL1 co-culture assays respectively ^25, 27^. We performed these assays at E10.5 as the YS is known to contain cells with lymphoid potential at this developmental stage ^53^. Here, we plated AGM or YS+VU cells without prior isolation of the traced or untraced fractions, and normalized the % of labeled B or T lymphocytes after culture to the initial % of tdTomato/ZsGreen within Kit+ cells. Labeled cells generated mature B and T lymphocytes in culture with significantly higher frequency when activated at E8.5, in both AGM and YS+VU (**Figure 5D,F; Figure S3C,D**). Interestingly, in these conditions myeloid cells were also generated more efficiently by cells labeled at E8.5 (**Figure 5E; Figure S3C**). These results suggest that while EMP emergence takes place from E7.5 onwards, progenitors displaying B- and T-lymphoid potential emerge from intra- and extra-embryonic HE between E8.5 and E9.5.

### Single cell transcriptomics coupled with HE lineage tracing identify distinct subsets of (pre-)HSCs in AGM and YS+VU

In order to determine the transcriptional identity of fetal-restricted HSPCs, we performed single cell transcriptomics of Ter119-cells isolated from *Cdh5-CreER^T2^::R26^tdTomato^* E10.5 YS+VU and AGM (**Figure S4A**). Cell clustering identified a mix of hematopoietic and non-hematopoietic populations (**Figure S4B**); therefore, we focused our further analyses on hemato-endothelial cells (**Figure S4C-D**). We identified 21 cell clusters (**Figure S4C**) that distributed along three main differentiation trajectories corresponding to myeloid, erythroid and Mk lineages (**Figure S4E**). Cluster #3, detected in both AGM and YS+VU, displayed a unique transcriptomic signature, expressing the hematopoietic genes *Myb*, *Kit*, *Runx1*, *Adgrg1, Flt3* together with many genes reported to identify HSCs or their immediate precursors, including *Hlf* ^33, 56, 59^, *Cd27* ^60, 61^, *Mecom* ^33, 62^, *Cd93* ^63^, *Hoxa7*, *Hoxa9* ^59, 64^ (**Figure S4F-G**). Based on their transcriptional signature, we identified these cells as (pre-)HSCs, as mature HSCs are nearly absent in the E10.5 embryo ^14, 65^. Interestingly, low levels of endothelial genes such as *Cdh5*, *Pecam1* and *Emcn* were detected in this cluster, suggesting an ongoing process of endothelial-to-hematopoietic transition (EHT) (**Figure S4F-G**). The (pre-)HSC signature in cluster #3 exhibited different levels of expression in AGM and YS+VU, suggesting heterogeneity (**Figure S4F-G**). To better evaluate the relationship between (pre-)HSC origin and molecular signature, we performed a new scRNA-Seq experiment on higher cell numbers. We pooled hemato-vascular CD31+ and/or Kit+ cells from either AGM or YS+VU and separated them into labeled and unlabeled cells (4-OHT at E8.5) (**Figure S5A-B**). A total of 44 clusters were isolated (**Figure S5C**). Interestingly, (pre-)HSCs recognized by their transcriptional signature distributed in two distinct clusters (**Figure 6A-B**). Cluster #1 was largely contributed by tdTomato-labeled AGM and unlabeled YS+VU cells; in contrast, tdTomato+ YS+VU cells made up >75% of cluster #2 (**Figure 6C-D**). Both clusters expressed *Cd27* (**Figure 6B, Figure S5D**). Differential gene expression (DGE) analysis between the two clusters highlighted genes involved in hematopoietic differentiation as well as metabolic and inflammatory genes (**Figure S5E-F**). To dissect (pre-)HSC heterogeneity at the transcriptional level, we restricted our further analysis to (pre-)HSC clusters (**Figure 6E**). Relative gene set comparison evidenced that the (pre-)HSC transcriptional signature was expressed at the highest level by labeled cells of the AGM (**Figure 6F**), that our whole-mount imaging preferentially locates within the intra-embryonic portion of the VU (**Figure 3**). In contrast, expression of genes characteristic of hematopoietic differentiation was higher in tdTomato+ cells of the YS+VU (**Figure 6G**). AGM and YS+VU (pre-)HSCs exhibited a distinct metabolic signature, with glycolysis prominent in AGM and OXPHOS genes expressed at a higher level in YS+VU (**Figure S6A**). Pro-inflammatory genes, previously suggested to play a role in AGM HSC development ^66-69^, showed heterogeneous expression. Interestingly, interferon response genes were highest in labelled cells of AGM, while Tnf, Nfkb, Tlr4 and innate immune response genes were more expressed in labelled cells of the YS+VU (**Figure S6B**). Heterogeneity between tdTomato+ AGM and YS+VU (pre-)HSCs was confirmed by DGE analysis, that highlighted higher expression of ribosomal genes in the latter, indicative of higher metabolic activity (**Figure S6C-D**)^70^. Genes involved in signaling pathways implied in HSC specification such as Notch, Shh and TGF-β ^71^ were highly expressed in tdTomato-AGM (pre-)HSCs, probably the most immature subset (**Figure S6E**). Taken together, our data shows evidence of molecular heterogeneity within the (pre-)HSC pool at the time of HSC emergence, and confirms at the transcriptomic level that 4-OHT at E8.5 in *Cdh5-CreER^T2^* mice labels the majority of E10.5 (pre-)HSCs, likely representing the precursors of fetal-restricted HSPCs. Our data also suggest that (pre-)HSCs in AGM and YS+VU represent cells at distinct stages of maturation, with those located in the YS+VU being primed for differentiation.

**Figure 6.**
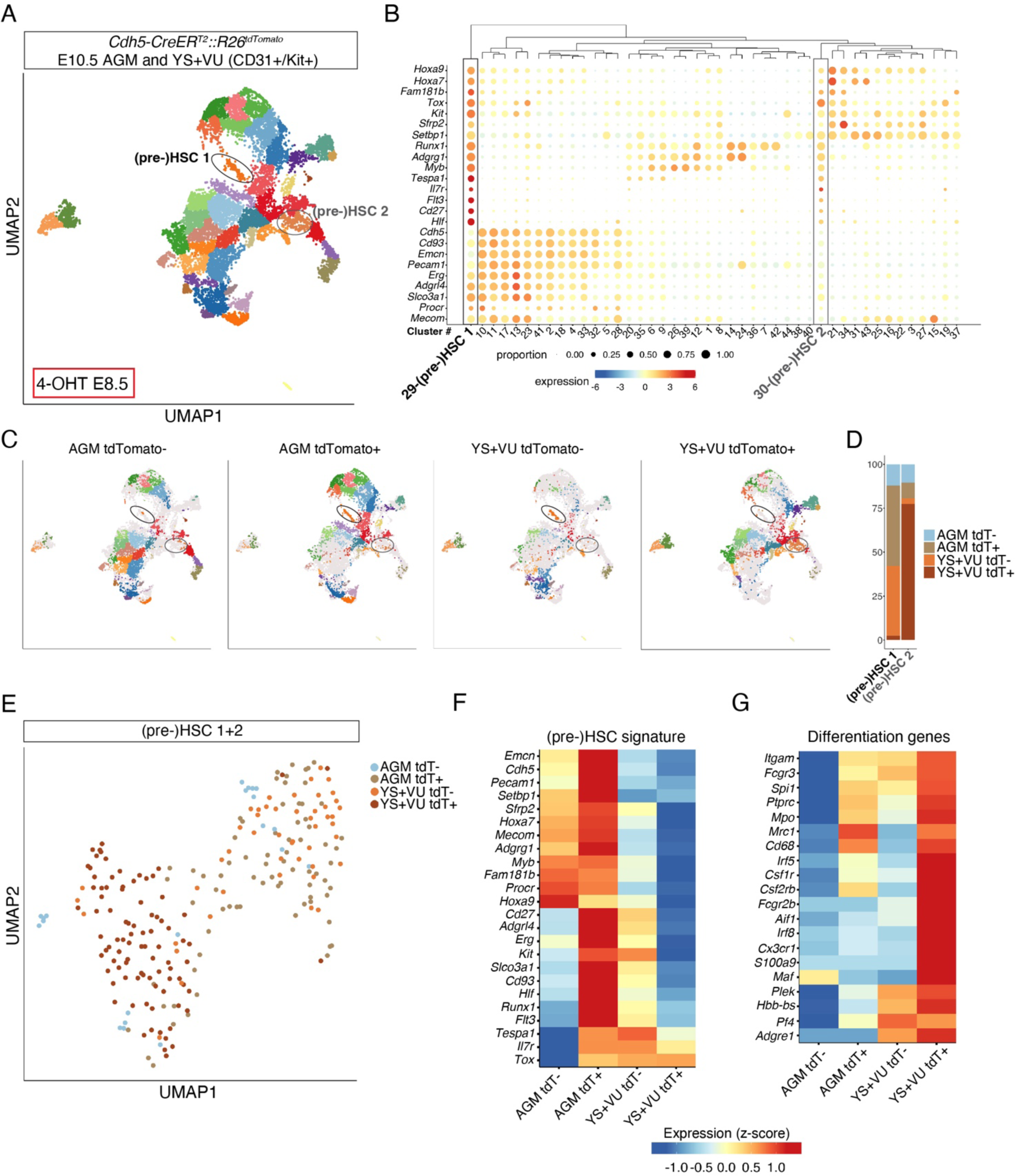
Lineage tracing and scRNA-Seq of E10.5 AGM, YS and extra-embryonic arteries identify distinct subsets of cells expressing a (pre-)HSC transcriptional signature. **(A)** Uniform Manifold Approximation and Projection (UMAP) of 26,180 cells (Ter119-Kit+/CD31+) isolated from AGM and YS+VU of E10.5 *Cdh5-CreER^T2^::R26^tdTomato^* embryos activated with 4-OHT at E8.5. AGM tdTomato+ (5,465), AGM tdTomato-(8,135), YS+VU tdTomato+ (4,445) and YS+VU tdTomato-(8,135) cells were separately sequenced (visual schematic of cell isolation strategy in S5A, gating strategy for cell isolation in S5B). Cells were isolated from 9 AGM and 14 YS+VU (1 litter of 9 embryos for AGM; 2 litters of 11 and 3 embryos for YS+VU). Cells are colored according to individual cluster identities, and (pre-)HSCs clusters #1 and #2 are circled (complete cell-type annotation is presented in Figure S5C). **(B)** Bubble plot showing the expression level of typical HSC and progenitor genes for each cell cluster in the whole dataset. Dot size indicates the percentage of cells expressing each gene, and dot color represents gene expression level. Black ((pre-)HSC 1) and gray ((pre-)HSC 2) boxes highlight clusters with a (pre-)HSC signature. **(C)** UMAP plots showing cell cluster distribution of AGM tdTomato-, AGM tdTomato+, YS+VU tdTomato- and YS+VU tdTomato+ samples. (pre-)HSC clusters 1 and 2 are circled. **(D)** Barplot showing the relative contribution of AGM tdTomato-, AGM tdTomato+, YS+VU tdTomato- and YS+VU tdTomato+ to (pre-)HSC clusters 1 and 2. **(E)** UMAP plot showing the subclustering of (pre-)HSC clusters 1 and 2. Dots represent (pre-)HSC cells colored by sample of origin. **(F)** Heatmap showing the relative expression levels of (pre-)HSC signature genes among AGM tdTomato-, AGM tdTomato+, YS+VU tdTomato- and YS+VU tdTomato+ cells within (pre-)HSC clusters #1 and #2. **(G)** Heatmaps showing the relative expression levels of hematopoietic differentiation genes among AGM tdTomato-, AGM tdTomato+, YS+VU tdTomato- and YS+VU tdTomato+ cells within (pre-)HSC clusters #1 and #2.

### Differentially labeled CD27+ Kit+ hematopoietic clusters selectively localize to vitelline and umbilical arteries

*Cd27* was one of the highly enriched genes in (pre-)HSCs. It was recently shown to be expressed in HSCs, type II pre-HSCs and lymphoid progenitors in E10.5-E11.5 AGM ^60, 61^. To assess whether CD27 would allow a better localization of pre-HSCs and progenitors with lymphoid potential within labeled cells, we performed whole-mount confocal imaging of E10.5 *Cdh5-CreER^T2^*embryos (**Figure 7A-E**). At this stage, we noticed that a higher percentage of cells within VU Kit+ cluster cells expressed CD27 as compared to DA or YS large clusters (**Figure 7B**), while clusters in the E9.5 YS did not express CD27 (**Figure S7A**), in agreement with the reported lack of CD27 expression in EMPs ^60^. Evaluation of labeling of Kit+ CD27+ cluster cells showed that E7.5 activation labeled a minority of those clusters in both AGM and YS (**Figure 7A,C,D,E**). In contrast, 4-OHT at E8.5 yielded significant labeling in the AGM (**Figure 7A,C**) and in YS arteries (**Figure 7D,E**). Strikingly, within the AGM, labeled Kit+ CD27+ clusters were detected in the VU but not in the DA (**Figure 7A,C**). These results strongly suggest that pre-HSCs and progenitors with lymphoid potential labeled with 4-OHT at E8.5 in *Cdh5-CreER^T2^* embryos specifically localize to VU, in both intra- and extra-embryonic regions.

**Figure 7.**
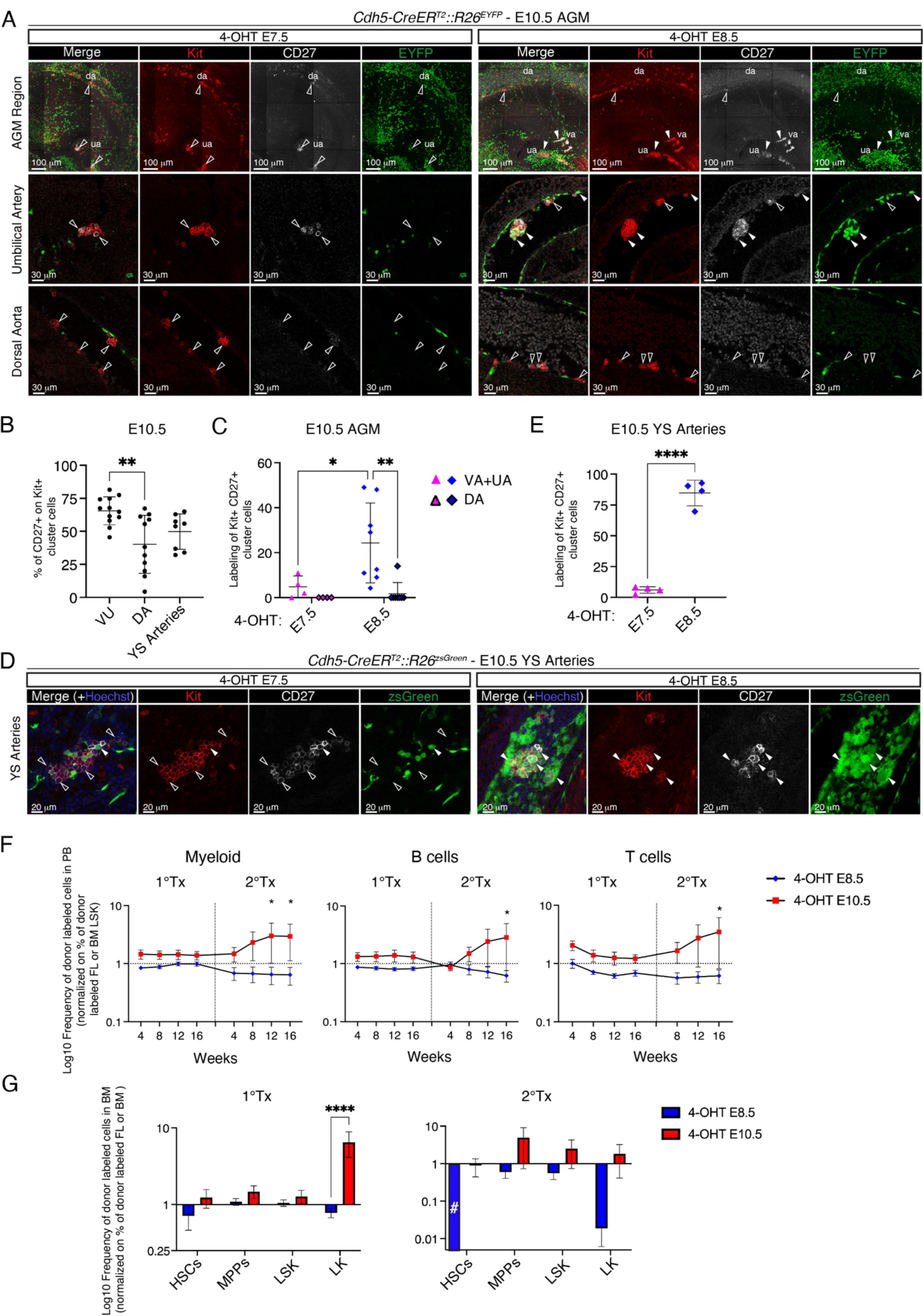
Localization of fetal-restricted HSPCs emergence in the VU arteries and assessment of their long-term engraftment potential. **(A)** Confocal WM-IF analysis of E10.5 *Cdh5-CreER^T2^::R26^EYFP^* AGM, activated with 4-OHT at E7.5 or E8.5. Upper panels show a 3D maximum intensity projection. Middle (UA) and lower (DA) panels show single 2.5 μm-thick optical slices. Arrowheads indicate examples of labeled (EYFP+; white arrowheads) or unlabeled (EYFP-; empty arrowheads) Kit+ CD27+ cells. 4-OHT E7.5 (n = 4) and 4-OHT E8.5 (n = 8) different YS were analyzed in 3 independent experiments. Scale bars: 100 μm (3D), 30 μm (slices). **(B)** Quantification of confocal WM-IF analysis shown in **(A-D)**, representing the percentage of CD27+ cells on the total of Kit+ cells. DA, VU (n=12) and YS arteries (n=8) were analyzed in 7 independent experiments. 3-9 images per sample were used. Error bars represent mean ± SD. **:p<0.01 (one-way ANOVA followed by Tukey’s multiple comparisons test). **(C)** Quantification of confocal WM-IF analysis in **(A)**. 4-OHT E7.5 (n= 4), and 4-OHT E8.5 (n=8) embryos were analyzed in 4 independent experiments. 5-15 images/embryo were used; 34 (4-OHT E7.5), 71 (4-OHT E8.5) different images were used. Error bars represent mean ± SD. *:p<0.05, **:p<0.01 (two-way ANOVA followed by Tukey’s multiple comparisons test). **(D)** Confocal WM-IF analysis of E10.5 *Cdh5-CreER^T2^::R26^zsGreen^* YS large arteries. Images show single 2.5 μm-thick optical slices. Arrowheads indicate examples of labeled (zsGreen+; white arrowheads) or unlabeled (zsGreen-; empty arrowheads) Kit+ CD27+ cells. 4-OHT E7.5 (n = 4) and 4-OHT E8.5 (n = 4) different YS were analyzed in 3 independent experiments. Scale bars: 20 μm. **(E)** Quantification of confocal WM-IF analysis in (**D**). 3-9 images/embryo were used. Error bars represent mean ± SD. ****p<0.0001 (two-tailed unpaired Student’s *t*-test). **(F)** Adult lethally irradiated mice were transplanted with E14.5 *Cdh5-CreER^T2^::R26^EYFP^* or *Cdh5-CreER^T2^::R26^tdTomato^* FL cells activated with 4-OHT at E8.5 or E10.5 (1° Tx) or adult BM cells from 1° TX (2° TX), (visual schematic of the experiment in S7B). Graphs show PB longitudinal analysis of transplanted mice, in which the frequency (Log10) of labeled (EYFP+ or tdTomato+) donor myeloid (CD45+ CD11b+), B cells (CD45+ B220+), T cells (CD45+ CD3e+) was normalized on the percentage of labeled LSKs (Lineage-Sca1+ Kit+) at the time of transplant. 4-OHT E8.5 1° Tx (n = 11), 4-OHT E10.5 1° Tx (n = 13), 4-OHT E8.5 2° Tx (n = 10), and 4-OHT E10.5 2° Tx (n = 11) recipient mice were analyzed in 2 independent experiments. Data is shown as mean ± SD. A comprehensive summary of PB analysis data in provided in Table S3 (1°Tx) and Table S4 (2°Tx). *p<0.05 (two-way ANOVA followed by Tukey’s multiple comparisons test). **(G)** BM analysis of 1° and 2° transplanted mice in (**F**), 16 weeks after transplant. Graphs show the Log 10 of frequency of labeled (EYFP+ or tdTomato+) donor HSCs (CD45.2+ Lin-Kit+ Sca1+ CD48-CD150+), MPPs (CD45.2+ Lin-Kit+ Sca1+ CD48+ CD150-), LSKs (CD45.2+ Lin-Kit+ Sca1+) and LKs (CD45.2+ Lin-Kit+ Sca1-), normalized on the initial labeling of each cell type in the donor tissue (FL for 1° Tx or BM for 2° TX). No labeled HSCs were found in 2°Tx recipients with 4-OHT at E8.5. Log10(0) is indicated with #. 4-OHT E8.5 1° Tx (n = 11), 4-OHT E10.5 1° Tx (n = 13), 4-OHT E8.5 2° Tx (n = 10), 4-OHT E10.5 2° Tx (n = 11) recipient mice were analyzed in 2 independent experiments. Data are shown as mean ± SD. A comprehensive summary of BM analysis data in provided in Table S4 (1°Tx) and Table S5 (2°Tx). ****p<0.0001 (two-way ANOVA followed by Tukey’s multiple comparisons test).

### Fetal-restricted HSPCs and adult-type HSCs exhibit distinct dynamics of engraftment

To establish whether fetal-restricted HSPCs could yield multi-lineage engraftment, we performed competitive transplants in which unfractionated cells from E14.5 *Cdh5-CreER^T2^::R26^tdTomato^*or *Cdh5-CreER^T2^::R26^EYFP^* FL pulsed with 4-OHT at E8.5 (fetal-restricted HSPCs) or E10.5 (adult-type HSCs) were transplanted into lethally irradiated syngeneic recipients (**Figure S7B**). The level of engraftment within donor-derived labeled fractions in PB was then followed over time and normalized to the percentage of labeled LSK in each of the donor samples (**Figure S7C**). Both E14.5 FL cells labeled at E8.5 and E10.5 exhibited multi-lineage engraftment potential in primary recipients (**Figure 7F, Figure S7D**). However, analysis of the progenitor compartment in BM of primary recipients at 16 weeks showed that only labeled cells pulsed at E10.5 could expand in the host BM niche (**Figure 7G, Figure S7E**). Next, we performed secondary transplantations (**Figure S7B**). In secondary recipients, the PB contribution of fetal-restricted HSPCs decreased, whereas that of HSCs significantly increased with time (**Figure 7F**). Upon terminal analysis, no labeled phenotypic HSCs from donor cells pulsed at E8.5 were detected in the BM of secondary recipients, while labeled progenitors originally pulsed at E10.5, including phenotypic HSCs, were still detected (**Figure 7G**). These results show that fetal-restricted HSPCs in a transplantation context behave in a similar way than in their physiological setting. Therefore, our data suggest that fetal-restricted HSPCs consist of a population of cells primed for differentiation, that does not contain HSCs with long-term multilineage engraftment potential.

## Discussion

Our results provide evidence of a wave of HSPCs exerting a major contribution to fetal lympho-myelopoiesis in mouse, which declines in postnatal life. This wave is sustained by progenitors emerging temporarily in between YS-derived EMPs and adult-type HSCs, the latter most probably originating primarily in the DA. Our genetic lineage tracing approach can label distinct waves of HE. We combined it with whole-mount imaging to demonstrate spatial segregation of the initial emergence of fetal-restricted HSPCs, localizing it within the HE of the VU. The first intra-embryonic hematopoietic clusters are generated in the portion of the VA most proximal to the DA at E9.5 ^12, 52, 53^. Though VU are known to harbor HSC precursors, this knowledge essentially relied on primary transplantation experiments ^10, 54^. Due to the lack of specific ways to identify and label VU-derived hematopoietic progenitors, their physiological role during development was unclear. Thus, this is the first report to trace their contribution in an unperturbed setting. The VA is directly connected to the UA at E9.5, and the emergence of fetal-restricted HSPCs at this stage coincides with the beginning of a dramatic vascular remodeling process, which eventually leads to the loss of direct connection between VA and UA and an extensive increase in size of the VA ^72, 73^. These events take place in the context of the establishment of a functional circulation ^74^. Although the VA can be already identified in the YS at E9.5 ^53^, at this stage we did not detect any differential labeling in Kit+ progenitors between 4-OHT activation at E7.5 or E8.5 in the YS. Thus, we hypothesize that the initial emergence of fetal-restricted HSPCs takes place in the intra-embryonic portion of the VA. The sudden presence of Kit+ clusters of differentially labeled progenitors in the YS arteries and UA at E10.5 shows that their appearance in these locations is an event taking place after E9.5, triggered either by *de novo* emergence from HE or by redistribution and proliferation of existing VU-derived progenitors. It is possible that VA remodeling and/or the onset of circulation play a functional role in the emergence of fetal-restricted HSPCs, as suggested by the smaller size of VA clusters and the complete absence of pro-HSCs in E9.5 *Ncx1-/-* embryos devoid of heartbeat ^52^.

Pulse-labeling of HE between E8.5 and E9.5, preferentially labeling hematopoietic clusters in the VU, allowed us to detect their extensive contribution to fetal lympho-myelopoiesis toward the end of gestation. Although with this activation mode we did observe variable labeling of phenotypic E14.5 FL HSCs, their labeling already decreased at E16.5, reaching a minimum in the adult. Recombination rates of the rest of the Lin-Kit+ progenitor compartment exhibited similar dynamics and correlated to the contribution to differentiated hematopoietic lineages. Therefore, our lineage tracing data, corroborated by single-cell transcriptomics and functional assays, provide evidence of a wave of fetal-restricted HSPCs poised for differentiation. Importantly, our transplantation experiments supported this notion by showing that fetal-restricted HSPCs are intrinsically programmed to a differentiation fate at the expense of self-renewal. Our data is consistent with previous reports suggesting functional heterogeneity within the E11.5 pre-HSC pool ^75^. Notably, pulse-labeling at E10.5 also recombined in a large fraction of E14.5 FL HSCs (75%), which gradually decreased to 54% at E16.5 and 40% in the adult BM (**Figure 1E-G**). Intriguingly, the extent of the decrease between labeled fetal and adult HSCs equals the percentage of lympho-myeloid cells (excluding macrophages) that were labeled in the E16.5 FL with this activation mode (∼30%; **Figure 2A**). This raises the possibility that 4-OHT at E10.5 may partially label fetal-restricted HSPCs in addition to adult-type HSCs, and suggests that fetal-restricted HSPCs may be virtually responsible for the entirety of lympho-myeloid contribution in the fetus. Our data are in agreement with recent results from Ganuza et al. showing that following an expansion phase between E12.5 and E14.5, most HSCs in the E14.5 FL preferentially undergo differentiation at the expense of self-renewal ^38^.

Although the existence of fetal restricted HSCs and MPPs had been suggested previously, the precise time and anatomical location of their emergence has never been investigated before. The first report to suggest the presence of developmentally restricted HSCs was based on *Flt3* lineage tracing ^37^; however, this study primarily relied on FL transplantation assays to probe their contribution, and offered little information on their origin and physiological role. Patel et al. also employed *Flt3* lineage tracing, but adopted an inducible strategy coupled with clonal barcoding to identify a population of eMPPs that arise early in embryogenesis and contribute to postnatal multi-lineage output ^34^. This study mainly focused on postnatal contribution and did not investigate in depth the emergence of eMPPs. Another recent report showed that hematopoietic progenitors in the FL are generated independently of adult-type HSCs and that *Mecom* levels, normally higher in the intra-embryonic region, can play a functional role in HSPC specification ^33, 48^. We found *Mecom* to be more expressed in AGM than YS+VU (pre-)HSCs (**Figure S4, Figure 6F**), consistent with an identity of the latter distinct from adult-type HSCs. The high expression of *Mecom* in labeled AGM pre-HSCs (**Figure 6F**) confirms that *Mecom*-expressing cells are predominantly located in the main arteries, including VU ^33^. Thus, Mecom expression levels may not only be important to specify adult HSCs, consistent with *EVI1^creERT2^* labeling phenotypic HSCs in the E14.5 FL ^33^, which our data suggest to be largely fetal-restricted (**Figure 1**). Overall, our data is in agreement with previous reports suggesting the existence of fetal-restricted HSPCs. Our lineage tracing strategy cannot discriminate between the separate contributions of individual progenitors (i.e. fetal HSCs or eMPPs), but rather identifies a distinct wave of HE at the source of fetal-restricted HSPCs, that we demonstrate to act as the main driver of fetal lympho-myelopoiesis. We show that fetal-restricted HSPCs phenotypically emerge as pre-HSCs from the HE of VU. We speculate that this HE wave contains the precursors of the pre-constituted FL hematopoietic hierarchy ^33, 48^, including fetal HSCs ^37, 38^ and eMPPs ^34^, whereas adult-type HSCs are mainly generated by a later HE of the DA.

There is some controversy in the literature regarding the identity of early TSPs. Although most evidence agrees that the initial seeding of the thymus takes place independently of HSCs ^30, 76-78^, it was also recently suggested, based on an inducible *Csf1r* lineage tracing approach, that the first TSP may be of HSC origin ^29^. Our data show that T cell progenitors in the E16.5 thymus largely originate from fetal-restricted HSPCs. The complementary levels of labeling seen in distinct TSP subpopulations, with higher levels of labeling in mature TSP with E8.5 activation and immature TSP labeled at a higher frequency with E10.5 activation, provide supporting evidence for the existence of two separate TSP waves with distinct origins ^45^. Thus, our results may help reconciling the discrepancies in the literature, as we do not formally exclude an HSC origin for the earliest TSPs. Nevertheless, our results imply that EMPs or adult-type HSCs are not responsible for the initial thymus seeding.

Whereas in layered hematopoiesis HSPC emergence is spatially segregated, all multi-lineage progenitors eventually converge in the FL ^1^. For a long time, the FL was thought to be an intermediate reservoir for HSC expansion after their initial generation in the AGM, in preparation to their lifelong residency in the BM. Several recent work revolutionized this concept and showed that FL is in fact a complex hub in which distinct waves of progenitors differentiate, expand and might even mature with mechanisms still largely unknown ^33, 34, 38, 63, 79-81^. Indeed, ours and others’ data imply that fetal-restricted HSPC and adult-type HSCs coexist in the FL, where they undergo different processes leading respectively to differentiation and self-renewal. Testing to what extent HSPC fate is intrinsically pre-determined, and what exactly is the role of the microenvironment will offer crucial insight. Based on our findings, it is tempting to speculate that before FL colonization, the emergence and transiting of fetal-restricted HSPC in the VU niche plays a functional role in determining their identity and fate. Metabolic ^52, 82^ and inflammatory factors ^66-69^ were shown to promote HSC development. Indeed, here we observed that these genes exhibit variable levels of expression in subsets of pre-HSCs of different origins (**Figures S5 and S6**). In particular, interferon signaling, more active in AGM-derived pre-HSCs (**Figure S6B**), is required for the switch between fetal and adult HSCs, beginning prior to birth ^47^. Therefore, it is possible that inflammatory cues are one of the main extrinsic factors required to specify adult-type HSC identity. Thus, it will be interesting to investigate what are the specific signals that regulate fetal-restricted HSPCs.

In summary, here we show that a wave of fetal-restricted HSPCs is the primary contributor to fetal lympho-myelopoiesis. We demonstrate that the initial emergence of fetal-restricted HSPCs takes place from HE localized within the VU, thus being segregated in space and time from that of adult-type HSCs. The *in vivo* fate of fetal-restricted HSPCs is mainly accomplished by differentiation during late gestation, followed by a sharp decline in postnatal life. Cells belonging to this wave are already transcriptionally distinct at the stage of pre-HSC. These data have important implications for the study of embryonic hematopoietic development and for the understanding of pediatric leukemias with pre-natal origins.

## Acknowledgements

We kindly acknowledge the contributions of Riccardo Gamberale, Anna Giovenzana, Giulia Di Meo and the personnel of the animal facilities at San Raffaele Scientific Institute and the University of Milan for support and technical help. We thank Matteo Iannacone for providing *R26^zsGreen^* mice and Andrea Ditadi for supplying OP9-Dl1 cells. Emanuele Azzoni was supported by a Fondazione Cariplo “Biomedical Research conducted by young researchers” grant n. 2018-0102, a Leukemia Research Foundation “New Investigator Blood Cancer Research Grant Program”, Award ID: 831382 and Cariplo Telethon Alliance grant n. GJC22013. Cristiana Barone was supported by Fondazione Umberto Veronesi. Alessandro Fantin was supported by the Fondazione Cariplo (2018-0298) and the Associazione Italiana per la Ricerca sul Cancro (AIRC) (22905). Silvia Brunelli was supported by the European Union’s Horizon 2020 research and innovation programme under Marie Skłodowska-Curie grant agreement No 860034. Filipa Timóteo-Ferreira was supported by the European Union’s Horizon 2020 research and innovation programme under Marie Skłodowska-Curie grant agreement No 860034 and Cariplo Telethon Alliance grant n. GJC22013. Rocco Piazza was supported by Associazione Italiana Ricerca sul Cancro (AIRC) IG-29341, Italian MUR Dipartimenti di Eccellenza 2023-2027 (l. 232/2016, art. 1, commi 314 - 337) and European Union - NextGenerationEU through the Italian Ministry of University and Research under PNRR - M4C2-I1.3 Project PE_00000019 “HEAL ITALIA”. Ana Cumano was supported by grants from Institut Pasteur, Institut National de la Santé et de la Recherche Médicale, Agence Nationale de la Recherche (grant Twothyme, grant EPI-DEV and grant DELSTAR), REVIVE Future Investment Program and Ligue Nationale contre le Cancer. Schematics were created in BioRender.com.

## Author Contributions

CBa conceived, designed and performed experiments, analyzed and interpreted data, made figures and wrote the manuscript; GQ, RO, FTF performed experiments, analyzed and interpreted data, edited the manuscript; ACa, AP, FSdS, MN, AD, AM, EDE, VB, GS, ACo performed experiments and analyzed data; MM, DdA, SS made scRNA-Seq libraries; SBo, SDM performed cell sorting; CDO provided technical support; CBi, RAP, RM, MFTRDB provided reagents and materials; ACu, SBr provided reagents and materials and interpreted data; AF performed experiments, interpreted data and provided reagents and materials; RP provided reagents and materials, analyzed and interpreted data and performed bioinformatic analysis; EA conceptualized, supervised and led the study, performed experiments, analyzed and interpreted data, made figures and wrote the manuscript. All authors read the manuscript and approved its final version.

## METHODS

### Mice and embryos

*Cdh5-CreER^T2^* ^39^, *Csf1r-iCre* ^83^, *R26^zsGreen^*, *R26^tdTomato^* ^84^ and *R26^EYFP^*^85^ transgenic mice were previously described and were genotyped according to reported protocols (further details and primers listed in **Table S1**). *R26^zsGreen^*, *R26^tdTomato^*or *R26^EYFP^* females aged 6 to 16 weeks were subjected to overnight timed matings with *Cdh5-CreER^T2^* or *Csf1r-iCre* males. Successful mating was judged by the presence of vaginal plugs the morning after, which was considered 0.5 days post conception (E0.5). Embryos were collected and dissected as previously described ^52, 86, 87^. E9.5-E11.5 embryos were carefully staged by counting of somite pairs; older embryos were staged by morphological criteria. For *Cdh5-CreER^T2^* fate mapping, a single dose of 37.5 mg/kg of 4-hydroxytamoxifen (4-OHT) dissolved in corn oil was delivered by intra-peritoneal (i.p.) injections to pregnant females at E7.5, E8.5 or E10.5. To counteract adverse effects of 4-OHT on pregnancies, 4-OHT solutions were supplemented with progesterone (18.75 mg/kg). All transgenic mouse lines were maintained on a CD45.2 C57BL/6 genetic background, with the exception of females used for timed matings in order to generate adult mice with 4-OHT activation during embryogenesis, which were instead of C57BL/6/FVB mixed background (F1). Mice were housed with free access to food and water at the San Raffaele Scientific Institute and University of Milan Institutional mouse facilities. All experiments were performed in accordance with experimental protocols approved by San Raffaele Scientific Institute and University of Milan Institutional Animal Care and Use Committees (IACUC).

### Flow cytometry analysis and cell sorting

Single cell suspensions were obtained from embryonic tissues (yolk sac, embryo/caudal part, fetal liver) by incubating for 15 min at 37 °C in calcium/magnesium free PBS supplemented with FBS 10%, Penicillin-Streptomycin 1%, EDTA 2mM and collagenase type I (Sigma) 0.12% (w/v), followed by mechanical dissociation by pipetting. Single cell suspension from fetal thymus was obtained by mechanical dissociation and passed through a 26-gauge needle. Peripheral blood (PB) samples were collected by tail vein bleeding using a scalpel; bone marrow (BM) was obtained by flushing long bones using a syringe and filtered in 40μm strainers. PB, BM and fetal liver (FL) samples were treated with the appropriate amount of RBC Lysis Buffer. Single cell suspensions were incubated with conjugated antibodies and processed for flow cytometry as previously described ^52, 86, 87^. A list of antibodies used for flow cytometry can be found in **Table S1**. Voltages, compensation and gates were set using unstained, single stained and fluorescence-minus-one (FMO) controls. Dead cells were excluded based on Hoechst 33258 (Hellobio) or 7-AAD (Sigma) incorporation. Flow cytometry data acquisition was carried out using a LSR Fortessa X-20 (BD) analyzer and BD FACSDiva software (version 8.0.2). Cell sorting was performed using a MoFLO Astrios cell sorter equipped with Summit software version 6.3 (both from Beckman Coulter), or a BD FACSAria II with BD FACSDiva software. An average sorting rate of 500-1000 events per second at a sorting pressure of 25 psi with a 100μm nozzle was maintained. Flow cytometric data was analyzed using FlowJo software version 10 (BD).

### OP9 co-cultures

Yolk sacs with vitelline/umbilical arteries (YS+VU) and caudal parts were dissected from E10.5 (32-36 sp) *Cdh5-CreER^T2^::R26^tdTomato^*or *Cdh5-CreER^T2^::R26^zsGreen^* concepti. Three YS+VU and caudal parts were each pooled and processed into single cell suspensions. One embryo equivalent (e.e.) from each sample was analyzed by flow cytometry to determine the percentage of labeling (tdTomato+ or zsGreen+) in Kit+ cells. One e.e. was plated on confluent OP9 or Delta-like 1-expressing OP9 (OP9-Dl1) ^88^ stromal cells in six-well plates in induction medium (αMEM, 10% FBS, supplemented with 10 ng/ml IL-7 and 10 ng/ml Flt3 ligand). At four days, exhausted medium was changed with fresh medium. At seven days, hematopoietic cells that grew in suspension were collected from each well, split 1 to 2, and plated on new OP9 and OP9-Dl1 confluent cells. The two resulting samples were considered a technical duplicate. Ten days after the start of the culture, non-adherent cells were collected and analyzed by flow cytometry.

### Colony-forming unit-culture (CFU-C) assays

YS+VU and caudal parts were dissected from E9.5 (21-27 sp) or E10.5 (32-36 sp) *Cdh5-CreER^T2^::R26^tdTomato^* or *Cdh5-CreER^T2^::R26^zsGreen^*or *Cdh5-CreER^T2^::R26^EYFP^* concepti. Three YS+VU and caudal parts were each pooled and processed into single cell suspensions. Labeled and unlabeled cells were isolated by flow cytometry. CFU-C assays were performed using Methocult M3434 (Stem Cell Technologies), as per manufacturer’s instructions. Cells were plated in duplicate dishes and cultured at 37°C, 5% CO_2_ in a humidified chamber. Colonies were scored after 7 days.

### Whole-mount immunofluorescence analysis and imaging

Whole-mount immunofluorescence (WM-IF) was performed as previously described ^52, 89^. Briefly, embryos and yolk sacs were dissected and fixed in a 4% paraformaldehyde solution in PBS for 30 minutes to 2 hours at 4°C. For E10.5 embryos, limb buds and body wall were removed before fixation to expose the aorta. Next, samples were treated with a permeabilizing-blocking solution (0.2% Triton X-100, 2% donkey serum, 2% FBS) and incubated overnight with primary antibodies. A second step of incubation with appropriate secondary antibodies was then carried out. Antibodies used for WM-IF are listed in **Table S1**. After staining, embryos were cleared in a benzyl alcohol-benzyl benzoate solution (BABB) and mounted as previously described ^89^. YS were cleared overnight in a 50% solution of glycerol in PBS at 4°C and then flat-mounted on Superfrost glass slides. Samples were imaged using a Zeiss 710 confocal microscope equipped with a LD LCI Plan-Apochromat 25x/0.8 Imm Corr DIC M27 objective or an EC Plan-Neofluar 40x/1.30 Oil DIC M27 objective. Confocal image acquisition was carried out using Zeiss Zen software version 2.3 SP1; image processing and analysis was carried out using IMARIS Viewer software version 9.7.2 (Bitplane), ImageJ/Fiji (versions 2.3.5-2.9.0) and Adobe Photoshop CC 2019.

### Single cell RNA Sequencing (scRNA-Seq) of E10.5 AGM and YS+VU

Aorta-gonad-mesonephros (AGM) and YS+VU from E10.5 *Cdh5-CreER^T2^::R26^tdTomato^* embryos (31-37 sp; 4-OHT at E7.5 or E8.5) were dissected and processed into single cell suspensions as described above. For the experiment in **Figure S4** (4OHT at E7.5), live Ter119-cells were isolated by FACS and divided into tdTomato+ and tdTomato-. For sample #1 (YS+VU), the same number of tdTomato+ and tdTomato-(1.5x10^4^+ 1.5x10^4^ cells) were mixed before preparing libraries. For sample #2 (AGM), tdTomato-cells were mixed in a 2:1 ratio to the tdTomato+ cells (2x10^4^+1x10^4^ cells). For the experiment in **Figure 6**, **Figure S5** and **S6** (4OHT at E8.5), live Ter119-CD31+/Kit+ cells were isolated by FACS and divided into tdTomato+ and tdTomato-. Labeled and unlabeled cells were analyzed separately.

Single-cell scRNA libraries were generated using a Chromium instrument (10x Genomics) with a Next GEM Single Cell 3’ kit. Libraries were quantified using a Qubit fluorometer (Thermo Fisher) and their profile was analyzed using a TapeStation instrument (Agilent). NGS sequences were generated using a Novaseq 6000 instrument (Illumina) with a target of 25000 reads/cell. Following multiplexing, raw fastq reads were processed using *cellranger* v6.1 and aligned against the mm10 mouse genome (GENCODE vM23/Ensembl 98) modified with the *mkref* tool to add an artificial ‘*tdTomato*’ chromosome. The associated genome annotation GTF file was modified accordingly. Filtered count matrices generated with *cellranger* were processed with *Seurat* v4.0 ^90^ package implemented in R (versions 3.2.3 – 4.2.1). Cells with genes count > 300 and < 8000 and fraction of mitochondrial reads < 0.20 were kept for downstream processing. For the dataset in **Figure S4**, after filtering, 4,325 YS+VU and 1,377 AGM cells were included for further analysis. The dataset in **Figure 6**, **Figure S5** and **S6** includes 26,180 cells after filtering (5,465 AGM tdTomato+, 8,135 AGM tdTomato-, 4,445 YS+VU tdTomato+ and 8,135 YS+VU tdTomato-). After converting individual matrices in Seurat objects via the Read10X function, data were normalized and transformed using the *SCTransform* Variance Stabilizing Transformation using the *glmGamPoi* method, while also regressing-out for feature count, percent of mitochondrial counts and cell phase. Data generated from both samples were subsequently integrated with a Canonical Correlation Analysis using the *PrepSCTIntegration*, *FindIntegrationAnchors* and *IntegrateData* commands, by using SCT as the normalization method. Dimensionality reduction of the integrated data was initially carried-on using Principal Component Analysis (PCA) and subsequently with Uniform Manifold Approximation and Projection (UMAP) algorithms, by retaining the first 30 principal components of the PCA. Clusters were identified with the Louvain algorithm; their number was selected using the Clustree tool ^91^ (version 0.4.3) by maximizing cluster stability. Individual cell types were identified by using SingleR ^92^ (version 1.0.1) as well as with manual data curation. Trajectory pseudotime analysis was performed using Monocle3 ^93^.

### Gene Set Enrichment Analysis (GSEA)

Gene Set Enrichment analysis (GSEA) was conducted using significantly differentially expressed genes (DEG) (p < 0.05) between (pre-)HSC2 and (pre-)HSC1, as well as between AGM tdTomato+ (pre-)HSCs and YS+VU tdTomato+ (pre-)HSCs. Analyses were carried out using R package ClusterProfiler (version 4.8.3) ^94^ querying the Gene Ontology (GO) database. The analysis was performed by setting the number of permutations to 10000, the minimum gene set to 3 and maximum to 800. GO terms were considered significant with a selected cutoff p-value of 0.05.

### In vivo transplantation

For transplantation experiments, syngeneic C57BL/6 (CD45.1) recipient mice were lethally irradiated (9 Gy, split dose) before intra-venous transplantation of 1x10^6^ unfractionated FL cells (primary transplantation) or 2x10^6^ adult BM cells (secondary transplantation) from *Cdh5-CreER^T2^::R26^TdTomato^*or *Cdh5-CreER^T2^::R26^EYFP^* mice (4-OHT at E8.5 or E10.5). Donor-derived chimerism and percentage of labeling (tdTomato+ or EYFP+) within donor cells was determined by flow cytometry in PB at 4, 8 and 12 weeks post transplantation, and in PB and BM at 16 weeks post transplantation. BM from primary and secondary transplanted mice was analyzed by flow cytometry to determine the percentage of labeling of donor hematopoietic stem/progenitor cells. A comprehensive summary of PB and BM transplantation analysis data in provided in **Table S4** (primary transplants) and **Table S5** (secondary transplants).

### Quantification and statistical analysis

Statistical analyses were performed using GraphPad Prism v10.2.1. No specific randomization method was used. Animals were allocated into experimental groups according to their genotype. No specific methods were used for blinding, but in general samples were collected from mice by one individual and then processed and analyzed by different individuals, at which time genotypes or experimental conditions of each sample were not known. To determine the level of significance, unpaired two-tailed Student *t*-test, one-way and two-way ANOVA followed by Tukey’s multiple comparisons test were used as indicated in figure legends. P<0.05 was considered statistically significant, and the level of significance is indicated by asterisks: * p < 0.05; ** p < 0.01; *** p < 0.001; **** p < 0.0001.

### Data availability

#### Lead contact

Further information and requests for resources and reagents should be directed to and will be fulfilled by the Lead Contact, Emanuele Azzoni

#### Materials availability

This study did not generate any new unique reagents.

#### Data and code availability

- scRNA-seq data of YS+VU and AGM from E10.5 *Cdh5-CreER^T2^::R26^tdTomato^* embryos (4-OHT at E7.5, accession numbers; SRR22189730 and SRR22189731; 4-OHT at E8.5, accession numbers: SRR28006358, SRR28006359, SRR28006360, SRR28006361) have been deposited in the Sequence Read Archive (SRA) data repository (NCBI) with the accession number BioProject ID: PRJNA898269. Data will be made publicly available upon publication.
- This study does not report original code.
- Any additional information required to reanalyze the data reported in this study is available from the lead contact upon reasonable request.

