## Supplementary Figures and Legends for "Hemogenic endothelium of the vitelline and umbilical arteries is the major contributor to mouse fetal lympho-myelopoiesis"

### SUPPLEMENTAL FIGURE TITLES AND LEGENDS

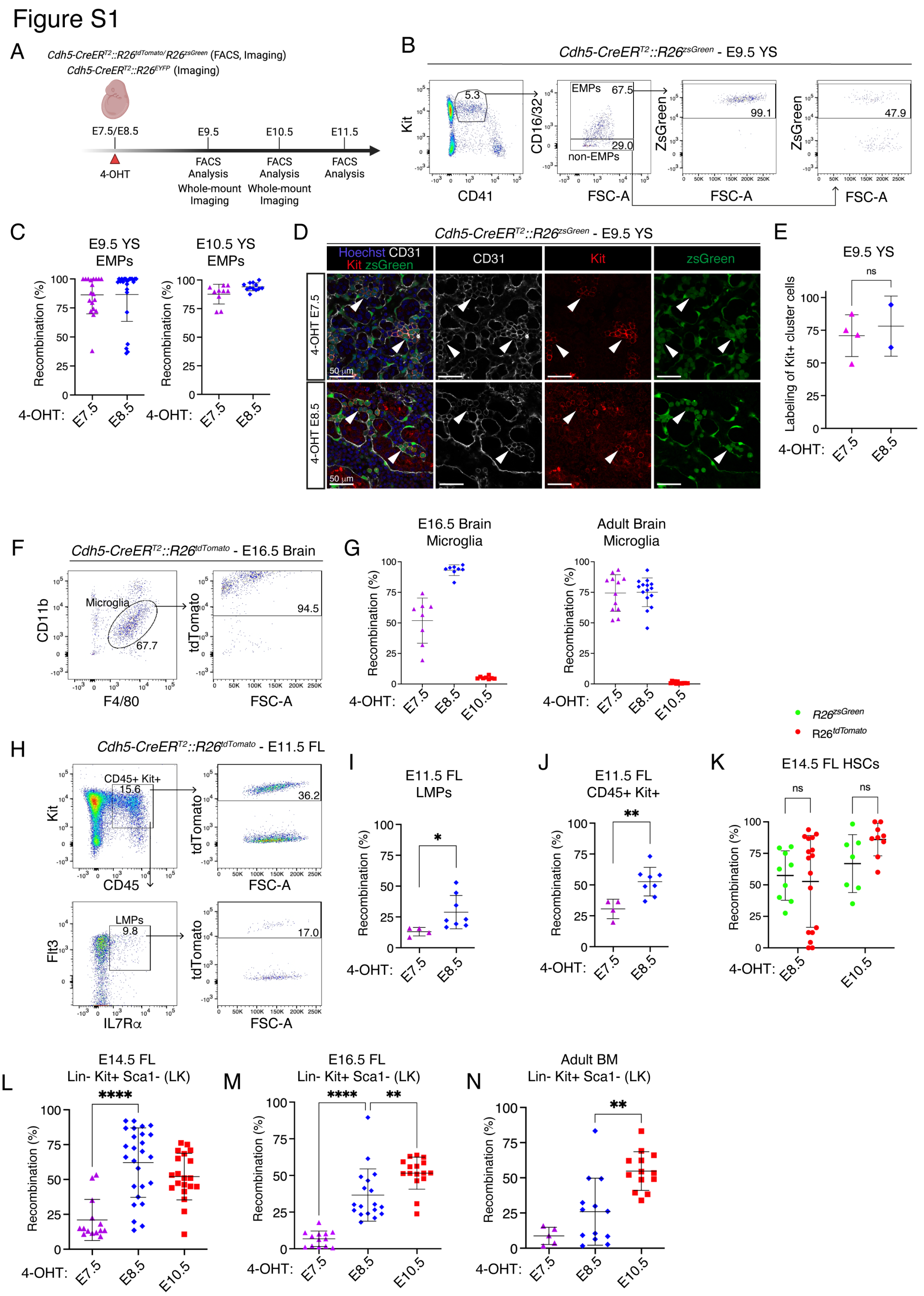

**Figure S1 (related to Figure 1). Lineage tracing analysis of EMPs, LMPs and LK progenitor cells in *Cdh5-CreER*$^{T2}$ embryos.**

**(A)** Visual schematic of lineage tracing experiments in E9.5, E10.5 and E11.5 embryos.

**(B)** Representative flow cytometric analysis of labeled EMPs (Ter119- $CD41^{lo}$ Kit+ CD16-32+) and non-EMPs (Ter119- $CD41^{lo}$ Kit+ CD16-32-) in E9.5 *Cdh5-CreER*$^{T2}$*::R26*$^{zsGreen}$ yolk sac (YS), activated with 4-OHT E7.5.

**(C)** Quantification of flow cytometric analysis in **(B)**. E9.5 4-OHT E7.5 (n = 19), E9.5 4-OHT E8.5 (n = 23), E10.5 4-OHT E7.5 (n = 10), E10.5 4-OHT E8.5 (n = 12) YS were analyzed individually in 5 independent experiments. Error bars represent mean ± SD.

**(D)** Confocal whole mount immunofluorescence (WM-IF) analysis of E9.5 *Cdh5-CreER*$^{T2}$*::R26*$^{zsGreen}$ YS. Arrowheads indicate CD31+ Kit+ hematopoietic cell clusters, labeled with 4-OHT at E7.5 (top) and E8.5 (bottom). 4-OHT E7.5 (n = 4), and 4-OHT E8.5 (n = 2) YS analyzed. Scale bar: 50 μm.

**(E)** Quantification of WM-IF analysis in **(D)**. Error bars represent mean ± SD. ns = non-significant (two-tailed unpaired Student's *t*-test).

**(F)** Representative flow cytometric analysis of brain microglia (CD45+ $CD11b^{low}$ F4/80+) in *Cdh5-CreER*$^{T2}$*::R26*$^{tdTomato}$ E16.5 embryos, activated with 4-OHT at E7.5, E8.5 (shown here) or E10.5. 4-OHT E7.5 (n = 8), (4-OHT E8.5 (n = 8), 4-OHT E10.5 (n = 8) were analyzed individually in 3 independent experiments.

**(G)** Quantification of flow cytometric analysis of labeled brain microglia as shown in **(F)** in *Cdh5-CreER*$^{T2}$*::R26*$^{tdTomato}$ E16.5 embryos (left) and 2-months-old adult mice (right), activated with 4-OHT at E7.5, E8.5 or E10.5. Replicates as shown. Error bars represent mean ± SD.

**(H)** Flow cytometric analysis of labeled LMPs (Lin- CD45+ Kit+ Flt3+ IL7Rα+) in E11.5 *Cdh5-CreER*$^{T2}$*::R26*$^{tdTomato}$ fetal liver (FL). 4-OHT E7.5 (n = 4), and 4-OHT E8.5 (n = 8) FL were analyzed individually in 2 independent experiments.

**(I)** Quantification of flow cytometric analysis of LMPs shown in **(H)**. Error bars represent mean ± SD. *p<0.05 (two-tailed unpaired Student's *t*-test).

**(J)** Quantification of flow cytometric analysis of labeled hematopoietic progenitor cells (CD45+ Kit+) shown in **(H)**. Error bars represent mean ± SD. ** p < 0.01, (two-tailed unpaired Student's *t*-test).

**(K)** Comparison of flow cytometric analysis of E14.5 FL HSCs as in **Figure 1**, shown separately for *Cdh5-CreER*$^{T2}$*::R26*$^{zsGreen}$ *and Cdh5-CreER*$^{T2}$*::R26*$^{tdTomato}$ embryos. Error bars represent mean ± SD.

**(L-N)** Quantification of flow cytometric analysis of labeled LK hematopoietic progenitor cells (Lineage- Kit+ Sca1-) related to **Figure 1D** in *Cdh5-CreER*$^{T2}$*::R26*$^{tdTomato}$ or *Cdh5-CreER*$^{T2}$*::R26*$^{zsGreen}$ E14.5 FL **(L)**, E16.5 FL **(M)** and adult BM (2-months-old) **(N)** activated with 4-OHT at E7.5, E8.5 or E10.5. . (E14.5 4-OHT at E7.5 (n = 14), E14.5 4-OHT at E8.5 (n = 26),

E14.5 4-OHT at E10.5 (n = 16), E16.5 4-OHT E7.5 (n = 13), E16.5 4-OHT E8.5 (n = 18), E16.5 4-OHT E10.5 (n = 17), 2 months 4-OHT E7.5 (n = 5), 2 months 4-OHT E8.5 (n = 13), 2 months 4-OHT E10.5 (n = 13) were analyzed individually in 17 independent experiments. Error bars represent mean ± SD. ** $p < 0.01$, **** $p < 0.0001$, (one-way ANOVA followed by Tukey's multiple comparisons test).

Figure S2

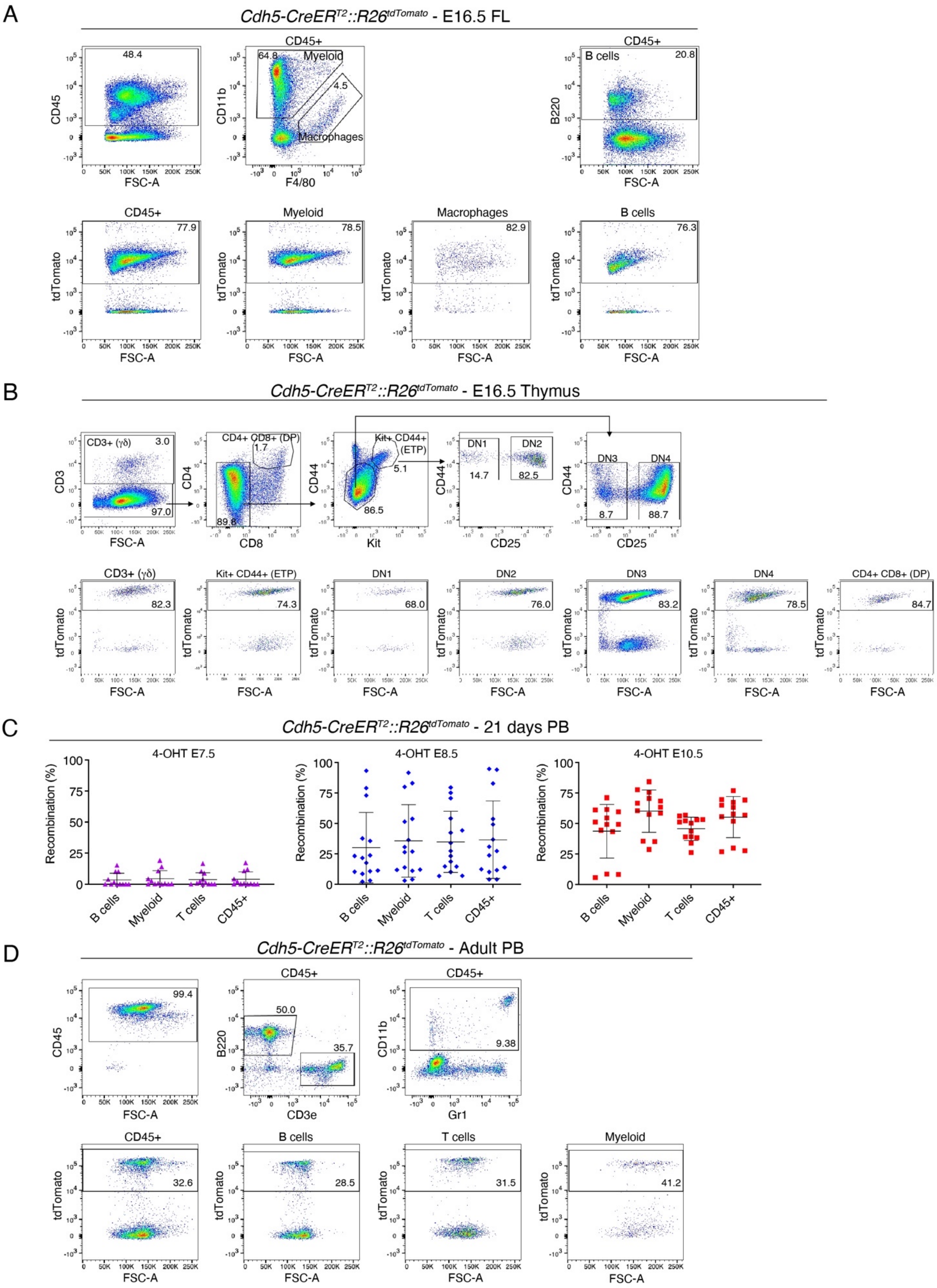
A
Cdh5-CreERT2::R26tdTomato - E16.5 FL
CD45+
Myeloid
Macrophages
B cells
48.4
64.8
4.5
20.8
77.9
78.5
82.9
76.3
CD45
CD11b
B220
tdTomato
FSC-A
F4/80
B
Cdh5-CreERT2::R26tdTomato - E16.5 Thymus
CD3+ (γδ)
CD4+ CD8+ (DP)
Kit+ CD44+ (ETP)
DN1
DN2
DN3
DN4
3.0
97.0
1.7
89.8
5.1
86.5
14.7
82.5
8.7
88.7
82.3
74.3
68.0
76.0
83.2
78.5
84.7
CD3
CD4
CD44
CD8
Kit
CD25
C
Cdh5-CreERT2::R26tdTomato - 21 days PB
4-OHT E7.5
4-OHT E8.5
4-OHT E10.5
Recombination (%)
B cells
Myeloid
T cells
CD45+
D
Cdh5-CreERT2::R26tdTomato - Adult PB
99.4
50.0
35.7
9.38
32.6
28.5
31.5
41.2
CD3e
Gr1

**Figure S2 (Related to Figure 2). Flow cytometric analysis of fetal and postnatal lympho-myeloid contribution in *Cdh5-CreER*$^{T2}$ mice.**

**(A)** Representative flow cytometric analysis of CD45+ (leukocytes), myeloid cells (CD45+ CD11b+), macrophages (CD45+ F4/80$^{hi}$), B cells (CD45+ B220+) in E16.5 *Cdh5-CreER*$^{T2}$*::R26*$^{tdTomato}$ FL, labeled with 4-OHT at E7.5, E8.5 (shown here) or E10.5. 4-OHT E7.5 (n = 21), 4-OHT E8.5 (n = 23), 4-OHT E10.5 (n= 8) FL were analyzed individually in 5 independent experiments. The corresponding quantification is shown in Figure 2A.

**(B)** Representative flow cytometric analysis of thymocytes in E16.5 *Cdh5-CreER*$^{T2}$*::R26*$^{tdTomato}$ or *Cdh5-CreER*$^{T2}$*::R26*$^{zsGreen}$ fetal thymus, labeled with 4-OHT at E7.5, E8.5 (shown here) or E10.5. 4-OHT E7.5 (n = 13), 4-OHT E8.5 (n = 18), 4-OHT E10.5 (n = 16) thymuses were individually analyzed in 4 independent experiments. Quantification shown in Figure 2B.

**(C)** Quantification of flow cytometric analysis of B cells (CD45+ B220+), myeloid cells (CD45+ CD11b+), T cells (CD45+ CD3e+), CD45+ (leukocytes) in juvenile 21 days old *Cdh5-CreER*$^{T2}$*::R26*$^{tdTomato}$ PB. 4-OHT E7.5 (n = 12), 4-OHT E8.5 (n = 15), and 4-OHT E10.5 (n = 13) mice were analyzed individually in 7 independent experiments. Error bars represent mean ± SD.

**(D)** Representative flow cytometric analysis of postnatal CD45+ (leukocytes), B cells (CD45+ B220+), T cells (CD45+ CD3e+), myeloid cells (CD45+ CD11b+) in *Cdh5-CreER*$^{T2}$*::R26*$^{tdTomato}$ PB labeled with 4-OHT at E7.5, E8.5 (2-months-old adult mice; shown here) or E10.5. 4-OHT E7.5 (n = 12), 4-OHT E8.5 (n = 13), and 4-OHT E10.5 (n = 13) mice were analyzed individually in 7 independent experiments. The corresponding quantification is shown in Figure 2C.

### Figure S3

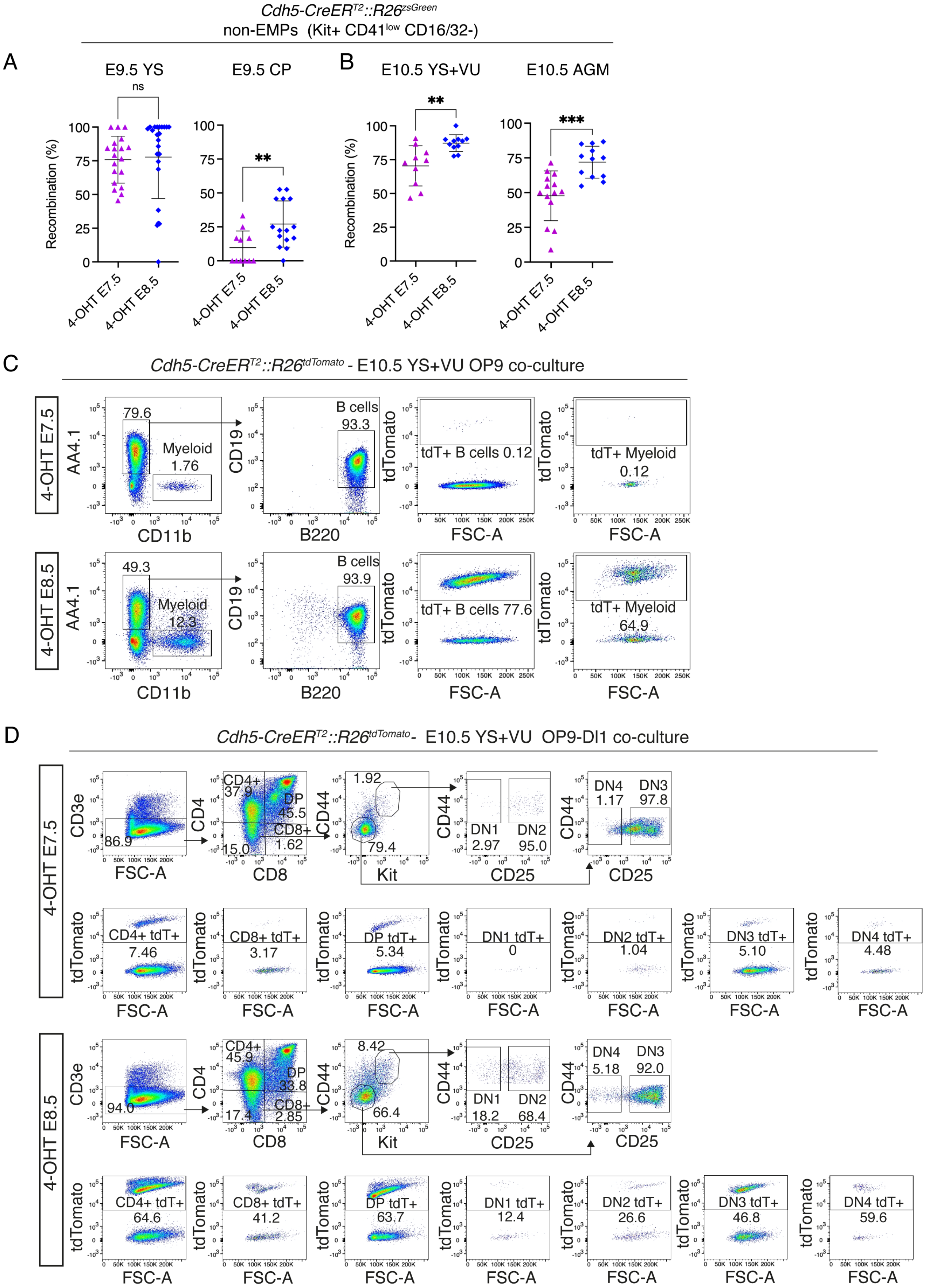

Cdh5-CreERT2::R26zsGreen
non-EMPs (Kit+ CD41low CD16/32-)
A
E9.5 YS
E9.5 CP
B
E10.5 YS+VU
E10.5 AGM
ns
**
**
***
Recombination (%)
4-OHT E7.5
4-OHT E8.5
C
Cdh5-CreERT2::R26tdTomato - E10.5 YS+VU OP9 co-culture
4-OHT E7.5
4-OHT E8.5
AA4.1
CD11b
Myeloid
CD19
B220
B cells
tdTomato
FSC-A
tdT+ B cells 0.12
tdT+ Myeloid
0.12
tdT+ B cells 77.6
64.9
D
Cdh5-CreERT2::R26tdTomato- E10.5 YS+VU OP9-Dl1 co-culture
CD3e
CD4
CD8
CD44
Kit
CD25
CD4+
DP
CD8+
DN1
DN2
DN3
DN4
CD4+ tdT+
CD8+ tdT+
DP tdT+
DN1 tdT+
DN2 tdT+
DN3 tdT+
DN4 tdT+

**Figure S3 (Related to Figure 5). Flow cytometric analysis of non-EMP hematopoietic progenitor cells and OP9/OP9-Dl1 co-cultures in *Cdh5-CreER$^{T2}$* embryos.**

**(A)** Quantification of flow cytometric analysis of non-EMP (Ter119- Kit+ CD41$^{low}$ CD16/32-) cells in *Cdh5-CreER$^{T2}$::R26$^{zsGreen}$* E9.5 YS (left) and caudal part (CP, right), labeled with 4-OHT at E7.5 or E8.5. Gates as shown in Figure S1B. 4-OHT E7.5 (n = 19), and 4-OHT E8.5 (n = 23) YS analyzed in 4 independent experiments. 4-OHT E7.5 (n = 11), and 4-OHT E8.5 (n = 15) CP analyzed in 3 independent experiments. Error bars represent mean ± SD. ns = non-significant; ** $p < 0.01$, (two-tailed unpaired Student's *t*-test).

**(B)** Quantification of flow cytometric analysis of non-EMP (Ter119- Kit+ CD41$^{low}$ CD16/32-) cells in *Cdh5-CreER$^{T2}$::R26$^{zsGreen}$* E10.5 YS including vitelline and umbilical arteries (VU), left and AGM, right, labeled with 4-OHT E7.5 or E8.5. Gates as shown in Figure S1B. 4-OHT E7.5 (n = 10), and 4-OHT E8.5 (n = 12) YS analyzed in 4 independent experiments. 4-OHT E7.5 (n = 14), and 4-OHT E8.5 (n = 12) AGM analyzed in 4 independent experiments. Error bars represent mean ± SD. ** $p < 0.01$; *** $p < 0.001$ (two-tailed unpaired Student's *t*-test).

**(C)** Flow cytometric analysis of B cell and Myeloid cells OP9 co-culture assay from E10.5 *Cdh5-CreER$^{T2}$::R26$^{tdTomato}$* YS and VU (YS+VU) activated with 4-OHT at E7.5 (top) or E8.5 (bottom). 4-OHT E7.5 B (n = 8), and 4-OHT E8.5 (n = 6) different samples were analyzed in 4 independent experiments. The corresponding quantification is shown in Figure 5D and E.

**(D)** Flow cytometric analysis of T cells and progenitors OP9-Dl1 co-culture assay from E10.5 *Cdh5-CreER$^{T2}$::R26$^{tdTomato}$* YS+VU activated with 4-OHT at E7.5 (top) or E8.5 (bottom). 4-OHT E7.5 (n = 7), and 4-OHT E8.5 (n = 6) different samples were analyzed in 4 independent experiments. The corresponding quantification is shown in Figure 5F.

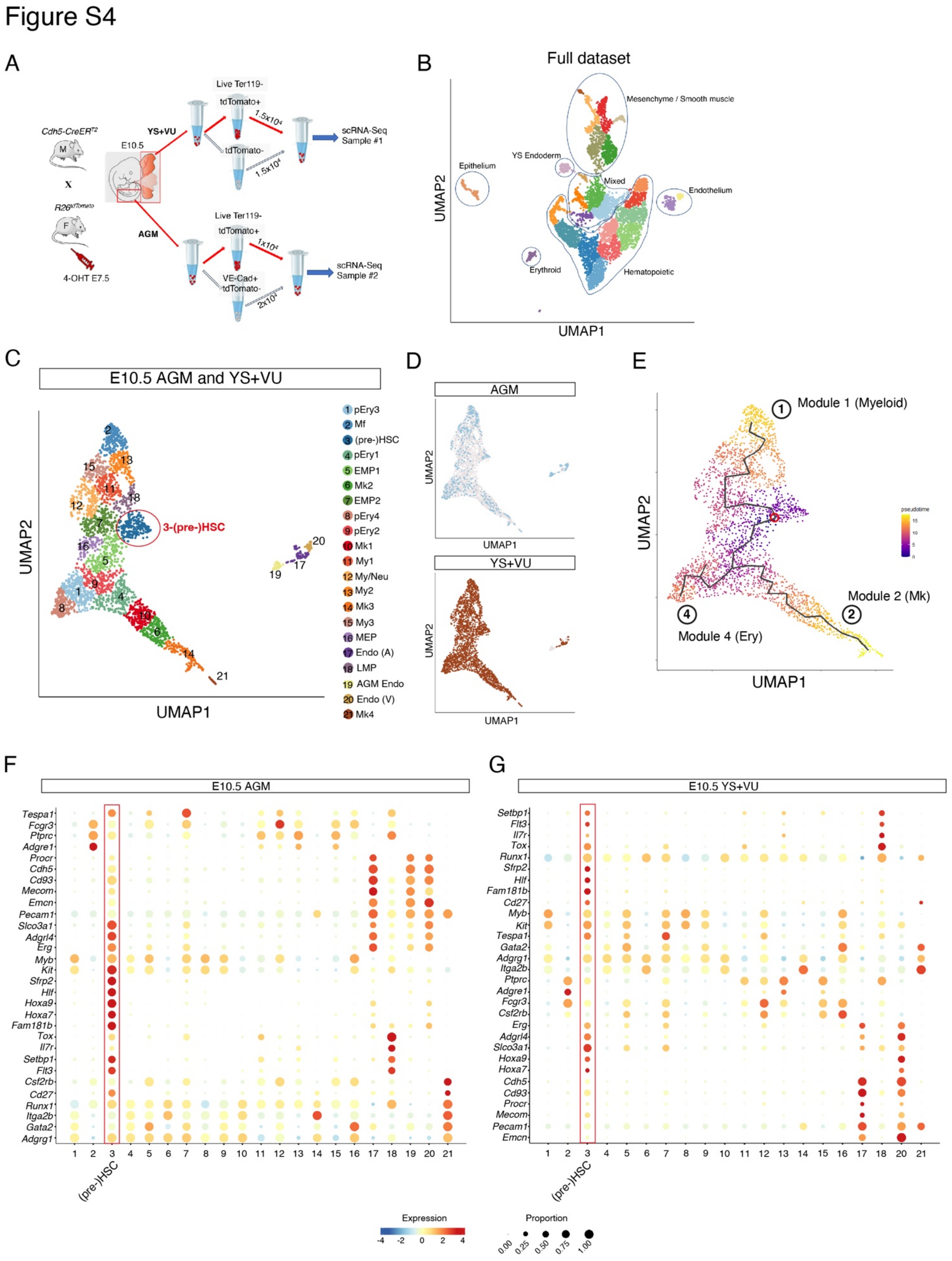

**Figure S4 (Related to Figure 6). scRNA-Seq identifies cells expressing a (pre-)HSC transcriptomic signature in E10.5 YS+VU and AGM.**

**(A)** Experimental schematic showing the cell isolation strategy used for single cell RNA sequencing (scRNA-Seq) of AGM and YS+VU from E10.5 *Cdh5-CreER*$^{T2}$*::R26*$^{tdTomato}$ embryos, (4-OHT activation at E7.5). 4-OHT at E7.5 was used since, based on our previous results, this activation differentially labels EMPs (present in tdTomato+ fraction) from fetal-restricted HSPCs (present in the tdTomato- fraction), thus allowing introduction of equal numbers of both fractions in the sequencing libraries.

**(B)** UMAP plot of 5,702 cells isolated from E10.5 Cdh5-*CreER*$^{T2}$*::R26*$^{tdTomato}$ AGM (1,377 cells) and YS+VU (4,325 cells) activated with 4-OHT at E7.5, that passed quality controls. Cells were isolated from 18 AGM and 18 YS+VU (2 different litters of 9 embryos each; cell isolation strategy shown in Figure S4A). Cells are colored according to cluster identity. Circled areas indicate different cell types annotated by gene expression markers.

**(C)** UMAP plot of the 3,206 hemato-endothelial cells (AGM, 602 cells; YS+VU, 2,604 cells) activated with 4-OHT at E7.5. Cells are colored according to cluster identity. This dataset was obtained excluding mesenchyme/smooth muscle, epithelium, mature erythroid and endoderm cells from the full dataset shown in Figure S4B. The (pre-)HSC cluster is circled in red.

**(D)** UMAP plots showing reciprocal distributions of AGM and YS+VU samples in the hemato-endothelial subset shown in (C).

**(E)** UMAP plot of the hemato-endothelial subset showing Monocle pseudotime analysis. Cells are colored according to their pseudotime score; the pseudotime trajectory is shown as a solid black line. Circled numbers correspond to main trajectory modules as identified by Monocle.

**(F)** Bubble plot showing expression of selected (pre-)HSC and progenitor genes for each cell cluster in the E10.5 AGM scRNA-Seq hemato-endothelial dataset. Dot size indicates the percentage of cells expressing each gene and color intensity represents gene expression level. Cluster 3 (red box) displays a (pre-)HSC signature.

**(G)** Bubble plot showing expression of selected (pre-)HSC and progenitor genes for each cell cluster in the E10.5 YS+VU. Dot size indicates percentage of cells expressing each gene and color intensity represents gene expression level. Cluster 3 (red box) displays a (pre-)HSC signature.

#### Figure S5

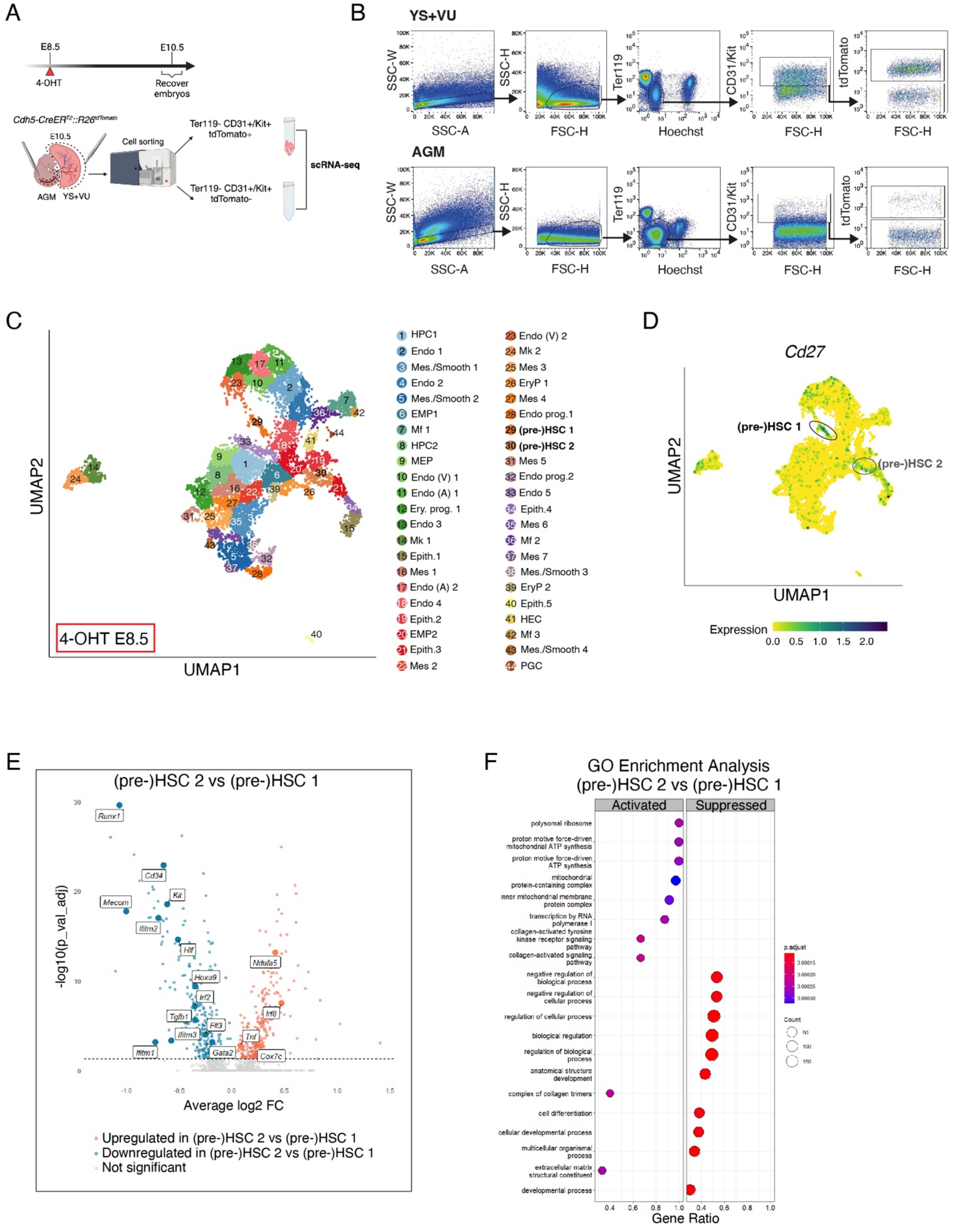

**Figure S5 (Related to Figure 6). scRNA-Seq identifies genes differentially expressed in distinct subsets of E10.5 AGM and YS+VU (pre-)HSCs.**

**(A)** Experimental schematic showing the cell isolation strategy used for scRNA-Seq on AGM and YS+VU from E10.5 *Cdh5-CreER*$^{T2}$*::R26*$^{tdTomato}$ embryos, with 4-OHT activation at E8.5. Hemato-endothelial (Ter119- CD31+/Kit+) cells were selected and tdTomato+ or tdTomato- fractions were separately sorted and sequenced from either AGM or YS+VU.

**(B)** Representative flow cytometric gating strategy for the isolation of live Ter119- CD31+/Kit+ tdTomato+ and tdTomato- for scRNA-Seq as represented in (A).

**(C)** UMAP plot showing the scRNA-Seq dataset obtained as in (A-B). Cells are colored according to cluster identity. Complete cell-type annotation was done according to the transcriptome.

**(D)** UMAP plot showing *Cd27* expression. Pre-HSC clusters 1 and 2 are circled.

**(E)** Volcano plot (Average log2 FC versus negative log of adjusted P value) used to visualize statistically significant gene expression changes (adjusted P value <0.05) between (pre-)HSC2 vs (pre)HSC1. Upregulated genes are labeled in red and downregulated genes labels in blue. Highlighted in labels are known genes related with hematopoiesis, inflammation and metabolic processes. A total number of 587 genes were differentially expressed (complete list in **Table S2**).

**(F)** Dotplot showing the GO biological process gene set enrichment analysis (GSEA) performed on the differentially expressed genes between (pre-)HSC2 vs (pre)HSC1. Dot size indicates the gene count and color intensity represents enrichment score (adjusted P value).

Figure S6

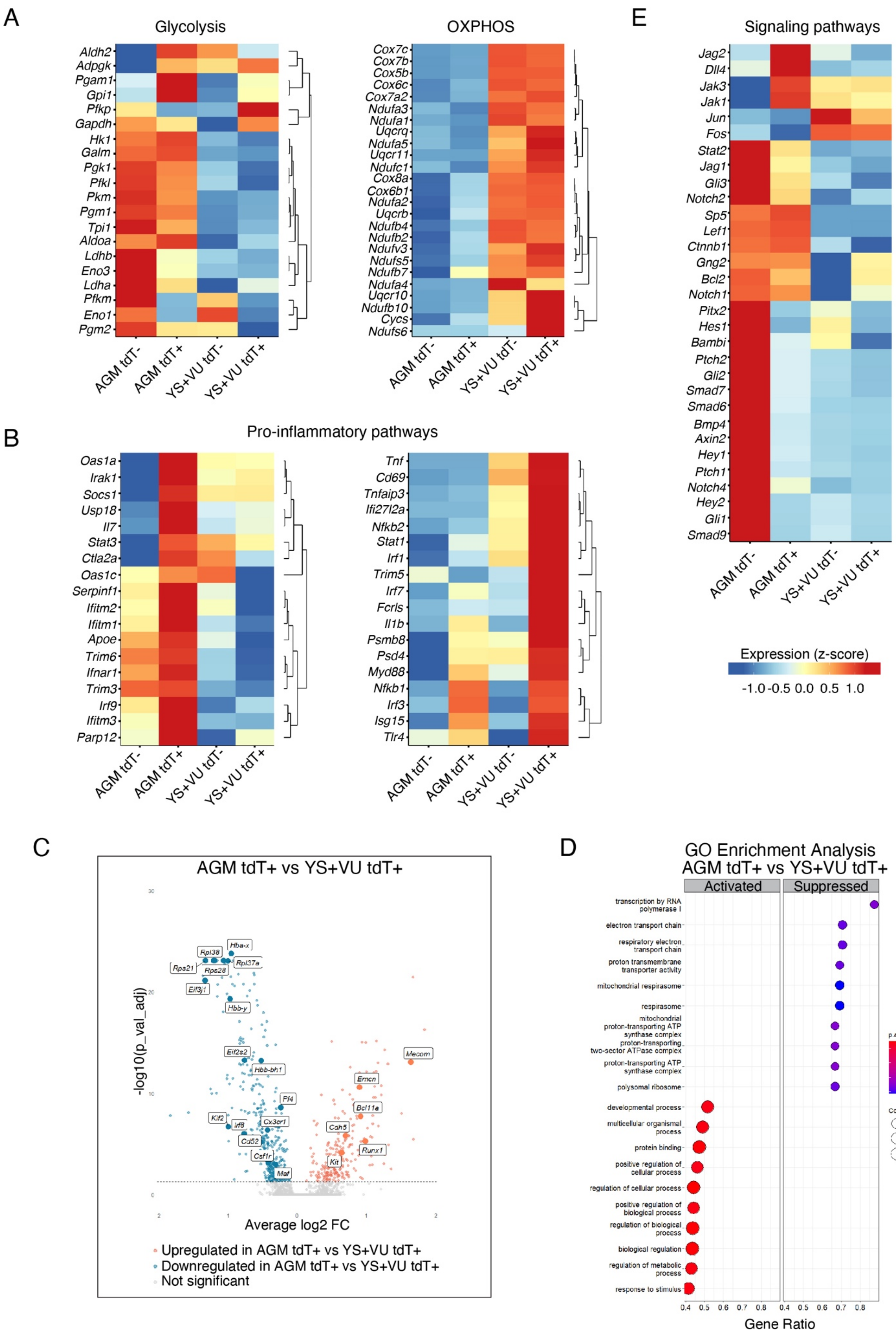
A
Glycolysis
OXPHOS
B
Pro-inflammatory pathways
C
AGM tdT+ vs YS+VU tdT+
-log10(p_val_adj)
Average log2 FC
Upregulated in AGM tdT+ vs YS+VU tdT+
Downregulated in AGM tdT+ vs YS+VU tdT+
Not significant
D
GO Enrichment Analysis
AGM tdT+ vs YS+VU tdT+
Activated
Suppressed
Gene Ratio
E
Signaling pathways
Expression (z-score)
-1.0 -0.5 0.0 0.5 1.0
AGM tdT-
AGM tdT+
YS+VU tdT-
YS+VU tdT+

**Figure S6 (Related to Figure 6). (Pre-)HSCs of distinct origin show different expression levels of genes involved in metabolism, inflammation and hematopoiesis.**

**(A)** Heatmap showing the relative expression levels of selected genes involved in the metabolic pathways of glycolysis (left) and oxidative phosphorylation (OXPHOS, right) among AGM tdTomato-, AGM tdTomato+, YS+VU tdTomato- and YS+VU tdTomato+ cells within (pre-)HSC clusters #1 and #2.

**(B)** Heatmap showing the relative expression levels of selected genes involved in pro-inflammatory pathways among AGM tdTomato-, AGM tdTomato+, YS+VU tdTomato- and YS+VU tdTomato+ cells within (pre-)HSC clusters #1 and #2.

**(C)** Volcano plot (Average log2 FC versus negative log of adjusted P value) used to visualize statistically significant gene expression changes (adjusted P value $<0.05$) between AGM tdTomato+ (pre-)HSCs vs YS+VU tdTomato+ (pre-)HSCs. Upregulated genes are labeled in red and downregulated genes labels in blue. Highlighted in labels are known hematopoietic and ribosomal genes. A total number of 698 genes were differentially expressed (complete list in **Table S3**).

**(D)** Dotplot showing the GO biological process gene set enrichment analysis (GSEA) performed on the differentially expressed genes between AGM tdTomato+ (pre-)HSCs vs YS+VU tdTomato+ (pre-)HSCs. Dot size indicates the gene count and color intensity represents enrichment score (adjusted P value).

**(E)** Heatmap showing the relative expression levels of selected genes involved in developmental and hematopoietic processes (e.g. Notch, BMP, Shh signaling pathways) among AGM tdTomato-, AGM tdTomato+, YS+VU tdTomato- and YS+VU tdTomato+ cells within (pre-)HSC clusters #1 and #2.

Figure S7

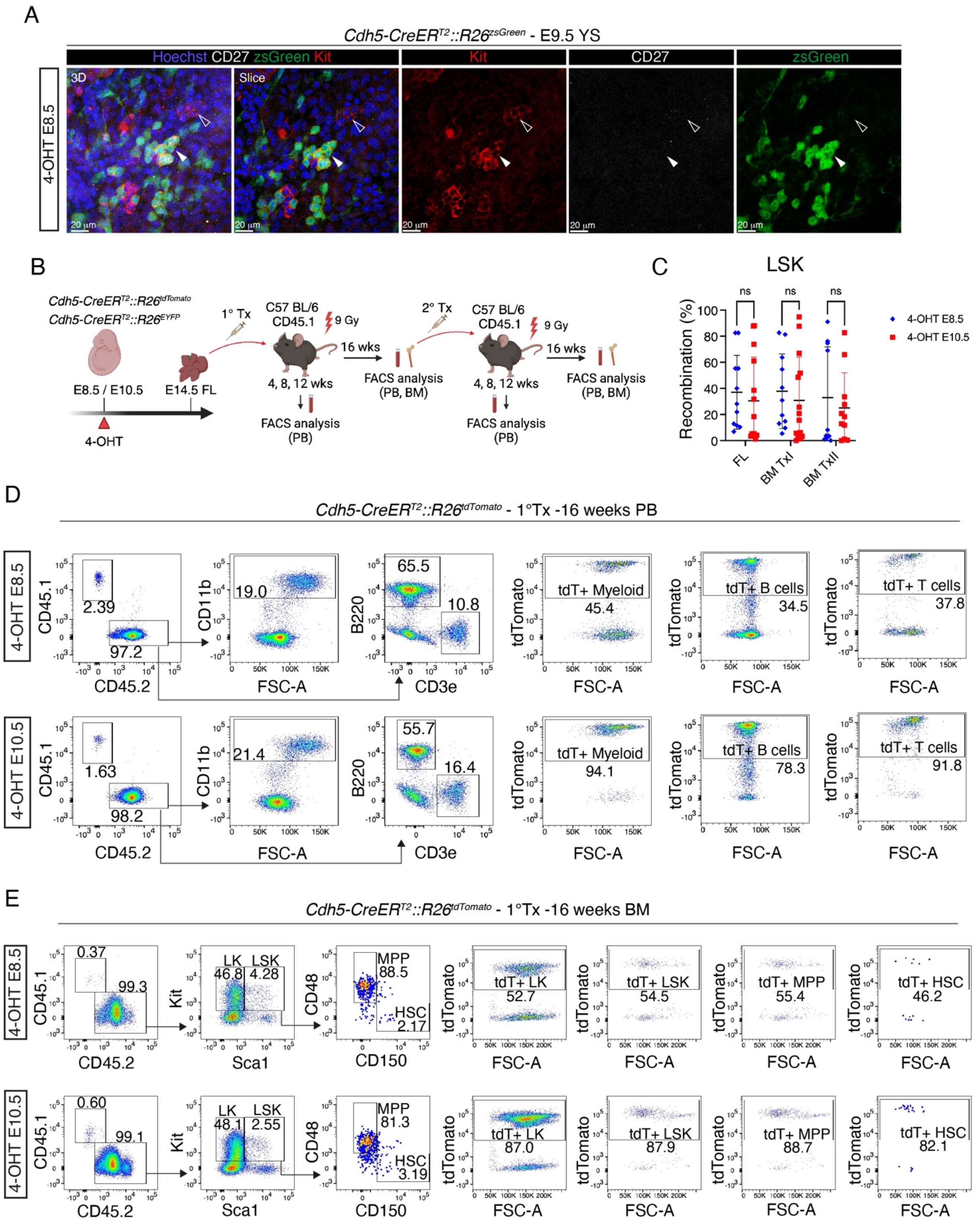

**Figure S7 (Related to Figure 7). CD27 expression in the E9.5 YS and flow cytometric analysis of E14.5 FL transplants.**

**(A)** Confocal WM-IF analysis of E9.5 *Cdh5-CreER*$^{T2}$*::R26*$^{zsGreen}$ YS (4-OHT E8.5) showing lack of CD27 expression within Kit+ clusters. Left panel shows maximum intensity 3D projection. Middle and right panels show single 2.5 µm-thick slices. Arrowheads indicate Kit+ CD27- clusters, labeled (zsGreen+; white arrowhead) or unlabeled (zsGreen-; empty arrowhead). A total number of 3 different YS were analyzed in 2 independent experiments. Scale bar: 20 µm.

**(B)** Visual schematic of E14.5 FL primary (1° TX) and secondary (2° TX) transplantation experiments.

**(C)** Quantification of flow cytometric analysis of LSK cells labeling in *Cdh5-CreER*$^{T2}$*::R26*$^{EYFP}$ *and Cdh5-CreER*$^{T2}$*::R26*$^{tdTomato}$ E14.5 transplanted FL and bone marrow (BM) of primary and secondary transplanted mice labeled with 4-OHT at E8.5 or E10.5. Number of replicates in Figure 7F. Error bars represent mean ± SD. ns = non-significant (two-way ANOVA followed by Tukey's multiple comparisons test).

**(D)** Representative flow cytometric analysis of PB from adult C57 BL/6 CD45.1 mice transplanted with *Cdh5-CreER*$^{T2}$*::R26*$^{tdTomato}$ E14.5 FL, activated with 4-OHT at E8.5 (top) or 4-OHT at E10.5 (bottom). FACS plots show hematopoietic populations and labeling percentages of donor myeloid cells, B cells and T cells. The corresponding quantification is shown in Figure 7F.

**(E)** Representative flow cytometric analysis of BM from adult C57 BL/6 CD45.1 mice transplanted with *Cdh5-CreER*$^{T2}$*::R26*$^{tdTomato}$ E14.5 FL, activated with 4-OHT at E8.5 (top) or 4-OHT at E10.5 (bottom). FACS plots show hematopoietic progenitor populations and labeling percentages of donor LKs, LSK, MPPs and HSCs. The corresponding quantification is shown in Figure 7G.

#### SUPPLEMENTAL TABLES

##### Table S1 - Reagents and Resources.

| REAGENT or RESOURCE | SOURCE | IDENTIFIER |
|---|---|---|
| Antibodies | | |
| Rat monoclonal anti-Ter119 APC-fire750 (clone TER-119) | BioLegend | Cat#116250; RRID: AB_2819833 |
| Rat monoclonal anti-CD117 (c-Kit) FITC (clone 2B8) | eBioscience | Ref: 11-1171-85; RRID: AB_465187 |
| Rat monoclonal anti-CD41 PE-Cy7 (clone eBioMWReg30) | eBioscience | Ref: 25-0411-82; RRID: AB_1234970 |
| Rat monoclonal anti-CD16/32 APC (clone 93) | BioLegend | Cat#101326; RRID: AB_1953273 |
| Rat monoclonal anti-CD150 PE-Cy7 (SLAM) (clone TC15-12F12.2) | BioLegend | Cat#115914; RRID: AB_439797 |
| Armenian hamster monoclonal anti-CD48 APC (clone HM48-1) | BioLegend | Cat#103412; RRID: AB_571997 |
| Rat monoclonal anti-Ly6A/E (Sca-1) Pacific Blue (clone E13-161.7) | BioLegend | Cat#122520; RRID: AB_2143237 |
| Rat monoclonal anti-CD117 (c-kit) APC-Cy7 (clone 2B8) | BioLegend | Cat#105826; RRID: AB_1626278 |
| Rat monoclonal anti-CD45 APC-eFluor 780 (clone 30-F11) | eBioscience | Ref: 47-0451-82; RRID: AB_1548781 |
| Rat monoclonal anti-CD117 (c-kit) PE-Cy7 (clone 2B8) | BioLegend | Cat#105814; RRID: AB_313223 |
| Rat monoclonal anti-CD135 Brilliant Violet 421 (clone A2F10) | BioLegend | Cat#135313; RRID: AB_2562338 |
| Rat monoclonal anti-CD127 (IL-7Ra) PE (clone A7R34) | BioLegend | Cat#135009; RRID: AB_1937252 |
| Rat monoclonal anti-Ter119 PE-Cy5 (clone TER-119) | BioLegend | Cat#116210; RRID: AB_313711 |
| Armenian hamster monoclonal anti-CD3e PE-Cy5 (clone 145-2C11) | BioLegend | Cat#100310; RRID: AB_312675 |
| Rat monoclonal anti-F4/80 PE-Cy5 (clone BM8) | BioLegend | Cat#123112; RRID: AB_893482 |
| Mouse monoclonal anti-NK1.1 PE-Cy5 (clone PK136) | BioLegend | Cat#108716; RRID: AB_493590 |
| Rat monoclonal anti-Ly6G/Ly6C (Gr1) PE-Cy5 (clone RB6-8C5) | BioLegend | Cat#108410; RRID: AB_313375 |
| Rat monoclonal anti-CD19 PE-Cy5 (clone 6D5) | BioLegend | Cat#115509; RRID: AB_313644 |
| Rat monoclonal anti-CD45R/B220 PE-Cy5 (clone RA3-6B2) | BioLegend | Cat#103210; RRID: AB_312995 |
| Rat monoclonal anti-CD45 PE (clone 30-F11) | BioLegend | Cat#103106; RRID: AB_312971 |
| Rat monoclonal anti-CD45R/B220 APC-Cy7 (clone RA3-6B2) | BioLegend | Cat#103224; RRID: AB_313007 |
| Rat monoclonal anti-CD11b PE-Cy7 (clone M1/70) | BioLegend | Cat#101216; RRID: AB_312799 |
| Rat monoclonal anti-Ly6G/Ly6C (Gr1) APC (clone RB6-8C5) | BioLegend | Cat#108412; RRID: AB_313377 |
| Mouse monoclonal anti-CD45.2 FITC (clone 104) | BioLegend | Cat#109806; RRID: AB_313443 |
| Mouse monoclonal anti-CD45.1 PE (clone A20) | BioLegend | Cat#110708; RRID: AB_313497 |

| | | |
|---|---|---|
| Rat monoclonal anti-CD93 (AA4.1) APC (clone AA4.1) | BioLegend | Cat#136510; RRID: AB_2275868 |
| Rat monoclonal anti-F4/80 APC (clone BM8) | BioLegend | Cat#123116; RRID: AB_893481 |
| Rat monoclonal anti-CD4 PE-Cy5 (clone RM4-5) | BioLegend | Cat#100514; RRID: AB_312717 |
| Rat monoclonal anti-CD8a PE-Cy7 (clone 53-6.7) | BioLegend | Cat#100714; RRID: AB_312753 |
| Rat monoclonal anti-CD25 PE-Cy7 (clone PC61) | BioLegend | Cat#102016; RRID: AB_312865 |
| Rat monoclonal anti-CD44 APC (clone IM7) | BioLegend | Cat#103012; RRID: AB_312963 |
| Mouse monoclonal anti-CD45.2 APC-Cy7 (clone 104) | BioLegend | Cat#109824; RRID: AB_830789 |
| Goat polyclonal anti-m/rCD31/PECAM1 | R&D systems | Cat#AF3628 |
| Rabbit polyclonal anti-GFP | Invitrogen | Cat#A11122; RRID: AB_221569 |
| Rat monoclonal anti-mouse CD117 (c-Kit) (clone 2B8) | eBioscience | Cat# 14-1171-82; RRID: AB_467433 |
| Rabbit polyclonal anti-RFP | Rockland | Cat# 600-401-379; |
| Armenian hamster monoclonal anti-mouse CD27 (clone LG-7F9) | eBioscience | Cat#14-0271-82; RRID: AB_467183 |
| Donkey polyclonal anti-goat Alexa Fluor Plus 647 | Invitrogen | Cat#A32849; RRID: AB_2762840 |
| Donkey polyclonal anti-rabbit Alexa Fluor Plus 488 | Invitrogen | Cat#A32790; RRID: AB_2762833 |
| Donkey polyclonal anti-rabbit Alexa Fluor Plus 555 | Invitrogen | Cat#A32794; RRID: AB_2762834 |
| Donkey polyclonal anti-rat Alexa Fluor 488 | Invitrogen | Cat#21208; RRID: AB_2535794 |
| Donkey polyclonal anti-rat CF568 | Biotium | Cat#20092; RRID: AB_ |
| Goat polyclonal anti-armenian hamster DyLight 649 (clone Poly4055) | BioLegend | Cat#405505; RRID: AB_1575122 |
| Rat anti-mouse CD16/CD32 antibody (Fc Block), clone 2.4G2 | BD Biosciences | Cat# 553142; RRID:AB_394657 |
| Rat Anti-Mouse CD44 BV510 (clone IM7) | BD Biosciences | Cat#563114; RRID: AB_2738011 |
| Hamster Anti-Mouse CD3e BUV395 (clone 145-2C11) | BD Biosciences | Cat#563565; RRID: AB_2738278 |
| Rat Anti-Mouse CD4 BV786 (clone RM4-5) | BD Biosciences | Cat#563727; RRID: AB_2728707 |
| Rat Anti-Mouse CD8a Pacific Blue (clone 53-6.7) | BioLegend | Cat#100725; RRID: AB_493425 |
| Rat Anti-Mouse CD43 Alexa Fluor 700 (clone S11) | BioLegend | Cat#143213; RRID: AB_2800660 |
| Biological samples | | |
| N/A | | |
| Chemicals, peptides, and recombinant proteins | | |
| 7-aminoactinomycin D (7-AAD) | BioLegend | Cat#420404 |
| Hoechst 33258 | Hellobio | Cat#HB0786 |
| Benzyl alcohol | Sigma-Aldrich | Cat#305197 |
| Benzyl benzoate | Sigma-Aldrich | Cat#68183 |
| MEM Alpha Medium | Gibco | Ref: 12000-063 |
| Sodium bicarbonate | Euroclone | Cat#ECM0980D |
| FBS HyClone | Cytiva | Cat#SH30071.03 |

| | | |
|---|---|---|
| Murine IL-7 | PeproTech | Cat#217-17 |
| Murine Flt3-Ligand | PeproTech | Cat#250-31L |
| Collagenase I | Merck | |
| 4-hydroxytamoxifen (≥98% Z isomer) | Sigma Aldrich | Cat#H6278 |
| Hot StarTaq Master Mix | QIAGEN | Cat#203445 |
| Progesterone | Sigma Aldrich | Cat#P0130 |
| Critical commercial assays | | |
| Methocult GF M3434 | STEMCELL Technologies | Cat# 03434 |
| Deposited data | | |
| Single cell RNA sequencing data of YS+VU and AGM from E10.5 *Cdh5-CreER*$^{T2}$*::R26*$^{tdTomato}$ embryos (4-OHT at E7.5) | This paper | BioProject ID: PRJNA898269 accession numbers; SRR22189730 and SRR22189731 |
| Single cell RNA sequencing data of YS+VU and AGM from E10.5 *Cdh5-CreER*$^{T2}$*::R26*$^{tdTomato}$ embryos (4-OHT at E8.5) | This paper | BioProject ID: PRJNA898269 accession numbers: SRR28006358, SRR28006359, SRR28006360, SRR28006361 |
| Experimental models: Cell lines | | |
| OP9 | Gift from Marella de Bruijn | N/A |
| OP9-Dl1 | Gift from Juan Carlos Zúniga-Pflücker | (Schmitt et al., 2004) |
| Experimental models: Organisms/strains | | |
| *Cdh5-CreER*$^{T2}$ | Gift from R.Adams | (Wang et al., 2010) |
| *R26*$^{zsGreen}$ | Gift from M.Iannacone | RRID:IMSR_JAX:007906 |
| *R26*$^{tdTomato}$ | The Jackson Laboratory | RRID:IMSR_JAX:007909 |
| *R26*$^{EYFP}$ | The Jackson Laboratory | RRID:IMSR_JAX:006148 |
| *Csf1r-iCre* | (Deng et al., 2010) | RRID:IMSR_JAX:021024 |
| B6.SJL-Ptprca Pepcb/BoyJ (B6 CD45.1) | Gift from L. Naldini | RRID:IMSR_JAX:002014 |
| Oligonucleotides | | |
| CreFW: TGATGGACATGTTCAGGGATC | Metabion | (Wang et al., 2010) |
| CreRV: CAGCCACCAGCTTGCATGA | Metabion | |
| zsGreen WT FW: AAGGGAGCTGCAGTGGAGTA | Metabion | (Madisen et al., 2010) |
| zsGreen WT RV: CCGAAAATCTGTGGGAAGTC | Metabion | |
| zsGreen TG FW: AACCAGAAGTGGCACCTGAC | Metabion | |
| zsGreen TG RV: GGCATTAAAGCAGCGTATCC | Metabion | |
| EYFP WT FW: CTGGCTTCTGAGGACCG | Metabion | (Srinivas et al., 2001) |
| EYFP WT RV: CAGGACAACGCCCACACA | Metabion | |
| EYFP TG FW: AGGGCGAGGAGCTGTTCA | Metabion | |
| EYFP TG RV: TGAAGTCGATGCCCTTCAG | Metabion | |
| tdTomato WT FW: AAGGGAGCTGCAGTGGAGTA | Metabion | (Madisen et al., 2010) |
| tdTomato WT RV: CCGAAAATCTGTGGGAAGTC | Metabion | |
| tdTomato TG FW: GGCATTAAAGCAGCGTATCC | Metabion | |
| tdTomato TG RV: CTGTTCCTGTACGGCATGG | Metabion | |
| Recombinant DNA | | |
| N/A | | |

| | | |
|---|---|---|
| Software and algorithms | | |
| Imaris (v. 9.7.2) | Bitplane | RRID: SCR_007370 |
| Zeiss Zen (v. 2.3 SP1) | Zeiss | https://www.zeiss.com/microscopy/int/products/microscope-software/zen.html |
| FlowJo (v. 10) | BD | RRID:SCR_008520 |
| GraphPad Prism (v. 9.4.1) | GraphPad Software | RRID:SCR_002798 |
| Photoshop CC 2019 | Adobe | RRID:SCR_014199 |
| Illustrator 2019 | Adobe | RRID:SCR_010279 |
| Microsoft Excel (v. 16.41) | Microsoft | RRID:SCR_016137 |
| Seurat (v. 4.0) | (Hao et al., 2021) | N/A |
| SCTransform | (Hafemeister and Satija, 2019) | N/A |
| R (R-3.2.3 – R-4.2.1) | The R Foundation | https://www.r-project.org |
| Louvain | (Blondel et al., 2008) | N/A |
| CellRanger (v 6.1) | 10x Genomics | https://support.10xgenomics.com |
| Clustree (v. 0.4.3) | (Zappia and Oshlack, 2018) | N/A |
| SingleR (v1.0.1) | (Aran et al., 2019) | N/A |
| Monocle3 | (Trapnell et al., 2014) | N/A |
| BioRender | BioRender.com | RRID:SCR_018361 |
| ImageJ (v. 1.54f) | ImageJ | https://imagej.net/ij/ |
| ShinyCell | (Liu et al., 2020) | http://shinycell1.ddnetbio.com |
| ClusterProfiler (v. 4.8.3) | (Wu et al., 2021) | N/A |
| Other | | |
| N/A | | |

**Table S2 (enclosed as a separate Excel file) – Differential gene expression analysis between (pre-)HSC2 and (pre)HSC1**

Differentially expressed gene (DEG) list between (pre-)HSC2 and (pre)HSC1 in the AGM and YS+VU E10.5 *Cdh5-CreER*$^{T2}$*::R26*$^{tdTomato}$ scRNA-Seq dataset with 4-OHT activation at E8.5. Statistically significant gene expression changes were considered as adjusted p value: <0.05.

**Table S3 (enclosed as a separate Excel file) – Differential gene expression analysis between AGM tdTomato+ (pre-)HSCs and YS+VU tdTomato+ (pre-)HSCs**

Differentially expressed gene (DEG) list between AGM tdTomato+ (pre-)HSCs vs YS+VU tdTomato+ (pre-)HSCs in the E10.5 *Cdh5-CreER*$^{T2}$*::R26*$^{tdTomato}$ scRNA-Seq dataset with 4-OHT activation at E8.5. Statistically significant gene expression changes were considered as adjusted p value: <0.05.

**Table S4 (enclosed as a separate Excel file) –E14.5 FL primary transplantation data**

Summary of peripheral blood and bone marrow analysis of adult C57 BL/6 CD45.1 mice transplanted with *Cdh5-CreER*$^{T2}$*::R26*$^{tdTomato}$ E14.5 FL, activated with 4-OHT at E8.5 or E10.5.

**Table S5 (enclosed as a separate Excel file) –E14.5 FL secondary transplantation data**

Summary of peripheral blood and bone marrow analysis of adult C57 BL/6 CD45.1 mice transplanted with BM from E14.5 primary transplant recipients.
